# How does fertilization impact the wild blueberry microbiome ?

**DOI:** 10.1101/2023.07.24.550391

**Authors:** Simon Morvan, Maxime C. Paré, Jean Lafond, Mohamed Hijri

## Abstract

Wild blueberries production is regarded as less intensive than other agricultural systems, although several agricultural practices are commonly implemented to increase crop yields and to mitigate pets and pathogen attacks. Fertilization, organic or mineral, is used to increase soil nutrient availability and improve fruit yield. Wild blueberry plants are also known to depend on their microbiome to overcome the lack of nutrient availability in the soil and their symbiosis with ericoid mycorrhizal fungi is thought to be crucial in that regard. As fertilization can alter crop microbial communities, our study aimed to measure the impact of this practice in a wild blueberry setting, focusing on the bacterial and fungal communities found in the roots and rhizosphere of *Vaccinium angustifolium* Ait., both at the time of application and more than year later during the harvest season. Our study indicates that fertilization, whether mineral or organic, has a minimal effect on microbial communities. One year after application, fertilization does not seem to have a negative repercussion on the ericoid mycorrhizal (ErM) fungal community as no significant differences were observed in terms of relative abundance on known and putative ErM taxa between the control and the two fertilizer treatments. The fact that fertilization is applied at a low rate could explain this absence of effect on the microbial communities. However, longer-term studies are needed to ensure that repeated fertilization does not cause any shifts in microbial communities that could be detrimental to the wild blueberry nutrition.

**Importance:** This study examines the impact of fertilization, whether organic or mineral, on microbial communities in the roots and rhizosphere of wild blueberry (*Vaccinium angustifolium* Ait.). both Fertilization is commonly used to enhance soil nutrient availability and improve fruit yield, but its effects on the plant’s microbiome, particularly on the ericoid mycorrhizal fungi (ErM), are not well understood. The samples were taken both during the pruning season, 3 months after the treatment, and one year later, during harvest season. The results suggest that fertilization has minimal impact on the microbial communities. One year after application, no significant differences were observed in the relative abundance of known and putative ErM taxa between the control group and the two fertilizer-treated groups. The low amount of fertilization applied could explain these results. However, longer-term research are needed to ensure that repeated fertilization does not lead to detrimental shifts in microbial communities affecting wild blueberry nutrition.

## Introduction

### Blueberry culture and agricultural practices

Wild blueberries, despite their name, are cultivated in Canada (Quebec and maritime provinces) and the United States of America (Maine). The vernacular name encompasses multiple species, mainly *Vaccinium angustifolium* Ait. and *Vaccinium myrtilloides* Michaux, that coexist in commercial blueberry fields (Yarborough, 2012). Contrary to the highbush blueberries culture (*V. corymbosum*) which resembles other fruit production with selected and planted cultivars, wild blueberries are grown from pre-existing stands that are usually found in abandoned farms or in the boreal forest understory. Once the land is cleared out of other plants and trees, wild blueberries propagate mainly through their rhizomes to progressively cover the entire field (Yarborough, 2012). This absence of sowing, transplanting or breeding explains the name of wild blueberries. However, similarly to other crops, producers manage their fields in order to maximize fruit yield. Among these agricultural practices are: pruning once every other year (Yarborough, 2004; Morvan *et al*., 2022b), pesticides applications to reduce problematic insects (Drummond and Groden, 2000; Collins and Drummond, 2004; Drummond *et al*., 2019), fungal diseases (Dedej et al., 2004; Esau *et al*., 2014; Hildebrand *et al*., 2016) and weeds (Jensen and Yarborough, 2004; Kennedy *et al*., 2010; Li *et al*., 2014) as well as fertilization (Eaton and Patriquin, 1988; Jeliazkova and Percival, 2003; Yarborough, 2004; Lafond, 2020).

### Fertilization effect on blueberry performance

Wild blueberries belong to the Ericaceae plant family along with cranberries (*Vaccinium macrocarpon*), rhododendrons (*Rhododendron spp.)* or heather (*Calluna vulgaris*). These plants are known to grow in acidic soils with low nutrient availability and high concentrations of trace metals. Essential nutrients are either contained in organic matter that cannot be directly absorbed by plants or bound to metals, mainly aluminum or iron in the case of phosphorus (Ohno and Severy, 2013). Wild blueberries are adapted to this environment and have relatively low nutrient requirements (Hall, 1978; Lafond, 2009). Nevertheless, studies have found that fertilizers could increase fruit productivity (Penney and McRae, 2000; Gagnon *et al*., 2003; Lafond, 2004; Starast *et al*., 2007; Maqbool *et al*., 2017). In Quebec, producers generally fertilize during the year following pruning using either mineral and/or organic amendments. Organic fertilizers have the advantage of slowly releasing nutrients, which could be more in phase with the wild blueberry nutrition requirements (Mallory and Smagula, 2012). However, organic fertilizers have the tendency to increase soil pH which can have a negative effect on ericoid mycorrhizal fungi, as they are adapted to acid soils (Haynes and Swift, 1985; Caspersen *et al*., 2016). Regarding their efficiency, Mallory and Smagula (2012) found no significant difference when they compared yields from fields fertilized with either mineral diammonium phosphate (DAP) or organic fertilizers (seafood waste compost and commercial organic fertilizers). One of the downsides of mineral fertilization, with mineral nitrogen in particular, is the ability of common blueberry weeds to uptake this element more efficiently. Therefore mineral nitrogen fertilization can generate weed proliferation if nothing is done to hamper their development, resulting in the opposite intended effect on blueberry yield (Penney and McRae, 2000; Marty *et al*., 2019).

### Wild blueberry microbial communities

The Ericaceae plant family forms a typical symbiosis within their roots with ericoid mycorrhizal (ErM) fungi (Leopold, 2016). These fungi produce a myriad of extracellular enzymes, allowing them to efficiently degrade multiple organic molecules present in the soil matrix (Kerley and Read, 1995; Cairney and Burke, 1998; Read *et al*., 2004; Martino *et al*., 2018). Nutrients released through this process are absorbed by those fungi and transferred to the Ericaceae hosts in exchange of carbohydrates produced by photosynthesis (Pearson and Read, 1973). Therefore, these fungal communities can play a major role in their host nutrition (Vohník *et al*., 2005) and studies have shown that inoculation with different ErM fungi increased plant growth and/or yield (Wei *et al*., 2016; Wazny *et al*., 2022). It has even been stated that the ability of wild blueberries to thrive in their harsh environment is due to this mycorrhizal symbiosis (Cairney and Meharg, 2003; Mitchell and Gibson, 2006). Furthermore, ErM fungi could also protect their host against trace metals toxicity through exclusion mechanisms (precipitation or fixation of metal ions outside of the plant cell, suppression of influx transporters, increased efflux of metal ions) or sequestration mechanisms (translocation of metal ions into subcellular organs where they are less toxic) (Perotto *et al*., 2002; Meharg, 2003; Mitchell and Gibson, 2006). Finally, other authors have shown that they increase their host tolerance to abiotic stress such as drought (Mu *et al*., 2021). Hence, a better understanding of these communities is important to minimize any disturbance of these beneficial fungi, and to try to harness their ecological service. In addition to the fungal community, bacteria are also regularly explored when studying plants’ rhizosphere and root microbiome (Adesemoye *et al*., 2009; Yang *et al*., 2009; Berendsen *et al*., 2012). To date, only a limited number of investigations have focused on the bacterial communities of wild blueberries or Ericaceae in general. However, bacteria could also be of importance, as studies have reported that Rhizobacteriales represented a large portion of wild blueberry rhizosphere bacterial communities (Morvan *et al*., 2020; Morvan *et al*., 2022). Furthermore, some of these Rhizobacteriales identified have previously been reported to be capable of nitrogen fixation, a number of which correlated to the wild blueberry leaf nitrogen content (Morvan *et al*., 2020; Morvan *et al*., 2022). Nevertheless further investigations are required to demonstrate nitrogen fixation by these bacteria in a wild blueberry soil setting. Additionally, the Rhizobacteriales order was also found to be abundant in multiple Ericaceae species (Timonen *et al*., 2017). Several other bacteria found in abundance in a previous study (Morvan *et al*., 2022) have previously been characterized and produce carbohydrate-active enzymes, enabling them to degrade multiple carbon sources, as is the case for several subdivisions of the Acidobacteriales order (Kielak *et al*., 2016), and Isosphaerales order (Kulichevskaya *et al*., 2016). Consequently, the bacterial community could also be of importance for the wild blueberry nutrition.

### Fertilization effect on microbial communities

Fertilization provides additional nutrients to crops, but can also strongly influence the plant’s microbiome (Weese *et al*., 2015; Li *et al*., 2020; Beltran-Garcia *et al*., 2021). This can depend on the species and growth stage of the plants, the conditions of cultivation (agricultural or unmanaged), the type and composition of the nutrients applied (mineral or organic), the application rate and the duration of fertilization (short or long term). Wild blueberries present a unique case, as the soils they grow in are acidic and they require relatively low amounts of nutrients. For instance, the recommended dose of ammonium sulfate in Quebec for wild blueberry is relatively low, between 30 and 70 kg N.ha-1 on a two-year cycle (Lafond and Ziadi, 2011; Lafond, 2020; Marty *et al*., 2019), whereas for spring wheat for example, the recommended N rate ranges from 90 to 120 kg N.ha-1 annually depending on soil texture (Nyiraneza *et al*., 2012). Although the mineral nitrogen fertilizers recommended for wild blueberry are ammonium-based (Lafond, 2009; Lafond, 2018), the lower amounts applied compared to other commercial crops might mitigate soil acidification and therefore lower the impact on the soil micro-organisms. To date, only one study has investigated the impact of fertilization on the wild blueberry microbiota. By testing different rates of N-P-K fertilizers (from 0 to 60 kg.ha-1) a study found a significant increase in ErM colonization when 35 kg.ha-1 of N in the form of urea was applied in the vegetative season (Jeliazkova and Percival, 2003).

The aim of our study was to document the effects of two types of fertilizers (organic and mineral forms) on fungal and bacterial root and rhizosphere microbiota. Since fertilizers are only applied during the vegetative growth stage, we wanted to investigate if any fertilizer effect on the fungal and bacterial communities could be observed a year after its application. Our hypotheses were: (1) the application of fertilizers at the beginning of the pruning growth stage will cause a shift in fungal and bacterial community composition, and if these shifts are observed, they will be stronger in the pruning growth stage than in the harvesting growth stage; (2) the mineral fertilisation will reduce the occurrence of ErM species as the plant will have readily accessible nutrients and are therefore less likely to require symbionts to sustain its needs; (3) the application of organic fertilization will increase the abundance of known ErM fungal species in the roots.

## Materials and Methods

### Experimental design

The experiment took place at the Bleuetière d’Enseignement et de Recherche located in Normandin, Quebec (48°49’40.2“N 72°39’36.9”W) in the North temperate zone (mean annual temperature: 0.9°C, annual precipitation: 871 mm (Government of Canada, 2021). This field was converted to blueberry production in 2005, and has been attributed for research purposes since 2016. The experiment took place in four adjacent fields, half containing the harvesting growth stage fields and the other the pruning growth stage fields. In each field, two blocks contained three 15×22m plot fertilization treatments (mineral, organic, none) resulting in four replicates for each fertilization and growth stage status for a total of 24 plots. Fertilization was applied during the pruning growth stage for all treated plots, before plant emergence, in early June. Therefore, in our study the pruning growth stage plots were fertilized in June 2019, and the harvesting growth stage plots were fertilized in June 2018 (Supplementary Figure 1). The mineral fertilization consisted in applying ammonium sulfate, triple superphosphate, potassium sulfate and borate (Millibore) at a rate of 50 kg of N.ha-1, 30 kg of P2O5.ha-1, 20 kg of K2O.ha-1 and 1.24 kg of B.ha-1 respectively. The organic fertilization provided the same concentration of N, P, K and B but using Actisol (5-3-2), a commercial natural hen manure fertilizer as well as the same dose of borate (Millibore).

### Soil chemical properties

Soil samples were extracted on the 29th of September 2019 in both the pruning and harvesting plots, using a soil corer with a 2.54 cm (1 inch) diameter at a rate of three pseudo-replicates per plot. The three pseudo-replicates were pooled into one sample once the organic layers (0 cm to 5 cm depth) and mineral layers (5 cm to 20 cm depth) were separated in the lab. Both layers were dried and sieved through a 2 mm mesh. Soil pH was measured using distilled water at a rate of 1:2 (Hendershot *et al*., 2007). Phosphorus, potassium, calcium and magnesium were extracted using Mehlich 3 solution (Ziadi and Tran, 2007). Phosphorus was quantified using colorimetry (Murphy and Riley, 1962), potassium with flame emission spectrophotometry while an atomic absorption spectrophotometer was used for calcium and magnesium (Perkin Elmer AAnalyst 300, Überlingen, Germany). Finally, nitrate, nitrite and ammonium were also measured by proceeding with a 1 M KCl extraction followed by a colorimetric dosage on a Lachat colorimeter (Hach Company, USA) using Quikchem® Method 12-107-06-2-A for ammonium, and Quikchem® Method 12-107-04-1-B for nitrate and nitrite (Maynard *et al*., 2007).

### Agricultural data

For the pruning year, leaves nutrient concentrations (N, P, K, Ca and Mg) were measured after a humid digestion of dried crushed leaves samples with a mixture of selenous acid and sulfuric acid - peroxyde (Isaac and Johnson, 1976; Lafond, 2009). For the harvesting year, these nutrients were measured in fruits instead of leaves, using the same method. Yields were also measured during the harvesting year using a mechanized harvester with a 1.5 m wide comb (rake) mounted on a Kubota F90 series tractor. The fruits were weighed directly in the field, but a sub-sample of 1 kg of blueberries was cleaned in the lab to remove debris (leaves, stems). The proportion accounted by the weight of debris was measured to apply a correction on the total fruit yield harvested.

### Microbial community analyses

#### Sampling

For the microbial community analysis, samples were collected between the 5^th^ and 14^th^ of August 2019. The sampling consisted in the collection of three, 10 cm wide and approximately 5 cm deep, clumps of organic soil per plot, containing blueberry rhizomes. Samples were placed in Ziploc bags, kept and transported into a cooler on ice until they could be placed at −20°C. Frozen samples were thawed to extract clumps of roots and soil from the original sample. The clumps obtained from the three pseudo-replicates were pooled into 50 mL Falcon tubes with pierced cap in order to freeze-dry the samples. The freeze-dried soil samples were then sieved through 1 mm mesh to remove coarse organic matter and pebbles. Roots and rhizomes were picked out of the sieve and placed in separate Ziploc bags. The resulting soil material, here forth named rhizospheric soil, contained thin root fragments that couldn’t be removed as well as organic matter. The rhizospheric soil was ground to homogenize the material to a fine powder. For the roots, we selected roots fragments with a diameter <1 mm in order to focus on the youngest roots and made sure to manually eliminate any visible soil particles.

#### DNA extraction and PCR amplification

The DNA extraction, amplification and sequencing protocol was similar to the one described in a previous study (Morvan *et al*., 2022). For the rhizosphere, 150 mg of soil were weighed to carry out the DNA extraction using the DNeasy PowerSoil kit (Qiagen, Toronto, ON) following the manufacturer’s protocol with the following modification: for cell lysis, samples were placed in TissueLyser II (Qiagen, Toronto, ON) instead of a vortex for 14 cycles of 45s each at speed 4.0. For the roots, we used 20 mg of ground roots to extract DNA using the DNeasy Plant kit (Qiagen, Toronto, ON) following the manufacturer’s instruction.

In addition to the DNA sample extraction, we included two negative controls consisting of DNA extraction of autoclaved and filtered water instead of either the roots or the rhizosphere soil. An additional PCR negative control was included by Genome Quebec. Two mock communities, fungal and bacterial, were also included in the sequencing procedure as positive controls. The fungal mock community was designed by Matthew G. Bakker and contains 19 fungal taxa (Bakker, 2018). In our experiment, we used the “even community” containing an equal amount of the 18S rRNA gene. The bacterial mock community contained 20 species (Supplementary Table 1) with equimolar counts (10^6^ copies/μL) of 16S rDNA genes (BEI Resources, USA). DNA extracts were stored at −20°C until they could be sent for amplification and sequencing by Genome Québec Innovation Center (Montréal, QC, Canada).

For the bacterial community, we targeted the V3-V4 region of 16S rDNA with the primers 341(F) and 805(R) resulting in an expected amplicon size of approximately 464 base-pair long (Mizrahi-Man *et al*., 2013). The sequencing platform (Centre d’expertise et de services Génome Québec, Montréal, QC) uses four versions of each primer (staggered primers) in order to increase base diversity in the Miseq flow cell. For the fungal community, we used the ITS3KYO(F) and ITS4(R) primers to target the ITS2 region located between the 5.8S and LSU region of the ribosomal RNA gene (Toju *et al*., 2012). The approximate amplicon length obtained with this pair of primers was 330 base-pair but this region is known to have varying length (Toju *et al*., 2012). Both sets of primers were coupled to CS1 (forward primers) and CS2 (reverse primers) tags that allow for barcoding. A second PCR was used to add a unique barcode per sample as well as i5 and i7 Illumina adapters that bind to the flow cell. Sequencing was conducted on an Illumina MiSeq using a paired-end 2 x 300 base-pair method (Illumina, San Diego, CA, USA).

#### Sequencing data processing

The sequencing data processing was identical to the one used in Morvan *et al*., (2022) as the sequences originated from the same sequencing run. Briefly, we used the DADA2 pipeline (Callahan *et al*., 2016a) in R (R Core Team, 2021) to process the raw fastq files. We processed the mock communities on their own in order to avoid any influence of our samples on the inferred mock ASVs. For both the fungal and bacterial datasets, we used cutadapt (Martin, 2011) in order to remove the primers, their complements, reverse and reverse complements by indicating the primers’ nucleotide sequences. For the fungal dataset, we proceeded with the filtering step using maxEE(2,2) and minLen(50) which removes sequences that are shorter than 50 nucleotides long. Regarding the bacterial dataset, we set truncLen to (270,240) and maxEE (2,2) based on a visual assessment of the quality profiles. After the different filtering step both bacterial and fungal dataset followed the same pipeline that used the default settings except those mentioned hereafter. In the learning error rates step, we used randomize = TRUE, in the sample inference step, we used “pseudo” as a pooling method and accordingly the “pooled” method for the bimera removal step. In order to add a taxonomy assignment to our inferred ASVs, we used the implemented naïve Bayesian RDP classifier with the *assignTaxonomy()* function. For the bacterial dataset, we used the SILVA reference database and two UNITE databases for the ITS sequences. Based on the taxonomy obtained, we removed non-bacterial ASVs as well as ASVs that were annotated as chloroplasts and mitochondria in the 16S dataset. The UNITE fungal reference database resulted in every ASVs to be labelled as Fungi but many sequences did not have an assigned Phylum. We compared this taxonomic assignment to a second one obtained by using UNITE’s eukaryotic reference database. Most of the unknown phyla fungi were labelled as non-fungal when using the eukaryotic reference database and were therefore removed in the ITS dataset. We then proceeded to further refine our datasets by removing singletons and doubletons (ASVs with a total abundance of 1 or 2) as they may be sequencing artefacts.

### Statistical analyses

All analyses were performed in R version 4.1.1 (R Core Team, 2021) and figures were generated using the ggPlot2 R package (Wickham, 2016). We looked at each growth stage separately as the harvesting and pruning blocks were not grouped together and had a different fertilization legacy.

#### Microbial data

To check if our sequencing depth allowed to capture most of the bacterial and fungal communities, we generated rarefaction curves using the *rarecurve()* function of the vegan R package (Oksanen *et al*., 2020). We used the phyloseq R package (McMurdie and Holmes, 2013) to facilitate data handling and to generate alpha diversity and beta diversity plots. We used Metacoder (Foster *et al*., 2017) for a more thorough analysis of the taxonomy of each dataset by plotting the ASV number and the relative abundance per taxa up to the genus level using the *heat_tree()* function.

#### Phylogenetic trees

In order to take into account the phylogenetic distance between the inferred ASVs, we built a phylogenetic trees following the method used in Callahan *et al*., 2016b. First, a sequence alignment was generated using AlignSeqs() from DECIPHER (Wright, 2016). We then used the phangorn package (Schliep, 2011) to compute a distance matrix under a JC69 substitution model using *dist.ml().* A neighbour-joining tree was then assembled onto which we fitted a generalized time-reversible with Gamma rate variation (GTR + G+I) model using *optim.pml().* The resulting trees were then added to their respective phyloseq objects.

#### Alpha and beta diversity

The Simpson and Shannon-Weaver alpha diversity indices were computed using the *plot_richness()* function of phyloseq. Statistical differences between the different fertilization treatments were checked using one-way ANOVA’s in a similar fashion than for soil and agricultural data (see above).

To assess the similarity of microbial communities, we relied on a beta diversity analysis using two different dissimilarity metrics. We took into account the compositional nature of sequencing data by choosing appropriate dissimilarity metrics (Gloor *et al*., 2017). First, we chose the Aitchison distance which consists in transforming the sequence abundances using a centered log ratio transformation before computing the Euclidean distance between sites (Quinn *et al*., 2018). Second, to take into account the phylogenetic distance between the sequences, we used the phylogenetic isometric log-ratio transformation PhILR before computing the Euclidean distance between sites (Silverman *et al*., 2017). Principal component analysis (PCA) ordinations based on both of these dissimilarity allowed to visualize the samples resemblance based on their microbial communities. To test the significance of the differences observed, we used vegan’s *adonis2()* function with 999 permutations and taking into account the experimental blocks. The homogeneity of dispersion assumption was checked using *betadisper()* and *permutest()* both in the vegan R package (Oksanen *et al*., 2020).

#### Representative taxa

Metacoder was used as a finer approach than the beta diversity analysis, to detect significant differences in each taxonomic rank between the burning intensities (Foster *et al*., 2017). The compare_groups() function was used after quantifying the per-taxon relative abundance using calc_taxon_abund(). This function computes a nonparametric Wilcoxon Rank Sum test to detect differences taxon abundance in different treatments. In order to take into account the multiple comparisons, we corrected the p-values of the Wilcoxon Rank Sum test with false discovery rate (FDR) correction using the p.adjust() function of the stats R package (R Core Team, 2021). After setting a threshold of 0.05 for the p-value, we used the *heat_tree_matrix()* to plot statistically different taxa based on their abundance using the log2 ratio of median proportions. Additionally, we also proceeded with an indicative species analysis to identify specific species representative of predefined groups of samples based on the species abundance and fidelity. Prior to this analysis, we agglomerated the ASVs at species level based on the obtained taxonomy. The significance of the results were obtained with permutation tests (9999 permutations) followed by an FDR correction to account for multiple testing. We used the multipatt() function of the indicspecies R package (de Cáceres and Legendre, 2009).

#### Redundancy analysis

We used constrained ordination in the form of distance-based redundancy analysis (db-RDA) to detect if the variation in the microbial datasets was linked to the variation of other variables such as fertilization, soil chemistry or agricultural variables, in a similar fashion as what was done in another study (Atashgahi *et al*., 2015). Prior to these analysis the ASVs were agglomerated using phyloseq function tax_glom if they had a similar species assigned. We used the distance matrices used for the beta diversity (Aitchison and PhILR distances) as the response variable and either the fertilization treatments, soil chemistry or agricultural variables as the explanatory variables. The fertilization treatments were converted to dummy variables (Legendre and Legendre, 2012). The soil chemistry and agricultural variables were standardized and we removed collinear variables prior to the db-RDA. The significance of the analysis was tested using permutation tests with the anova.cca() function of the vegan R package (Oksanen *et al*., 2020).

### Accession number

The microbial sequences have been deposited in the NCBI GenBank Sequence Read Archive under the accession number PRJNA995568: https://www.ncbi.nlm.nih.gov/bioproject/PRJNA995568.

## Results

### Microbial community overview

For bacteria, raw sequencing data resulted with an average of 87 000 reads per sample in both the rhizosphere and root samples. In the rhizosphere, the DADA2 pipeline filtered out almost 80% of the initial reads, while more than half of these initial reads were kept for the root samples. Despite this difference, approximately the same number of ASVs per sample were obtained in both compartments (Table 1). Additionally, 16 Archaeas, 69 Chloroplasts and 39 Mitochondrias ASVs were removed, along with 1128 ASVs that were only present once or twice across the dataset. After these steps, and excluding the control samples, 5869 bacterial ASVs summing up to 1 008 592 reads were spread across 48 samples. Despite the severe data loss during this filtering process, the rarefaction curves obtained for the bacterial community show clear inflexion points, and sometimes even reach a plateau. This indicates that much of the diversity has been captured, despite data loss during the filtering step (Supplementary Figure 2). Looking at the negative controls, we found five common ASVs in the rhizosphere extraction blank and in our dataset as well as 9 ASVs common in the root extraction blank and our dataset. Although they could originate from contamination, we did not exclude them from our dataset and treated these ASVs with caution in downstream analysis. The bacterial mock community identified 18 out of the 20 species present in the mock, and the two remaining were correctly assigned to the genus level. We did not observe any abundance bias towards a particular taxonomic lineage, though the mock lacked both Acidobacteriota and Planctomycetota representative taxa while these two phyla represent a significant proportion of the community found in our dataset.

**Table 1.** Bioinformatic steps.

|  | Bacteria |  | Fungi |  |
| --- | --- | --- | --- | --- |
|  | Rhizosphere | Root | Rhizosphere | Root |
| Initial read number | 87 698 ± 13 678 | 86 958 ± 10 534 | 66 848 ± 10 508 | 67 764 ± 14 224* |
| DADA2 processed | 19 658 ± 6 329 | 47 913 ± 8646 | 29412 ± 5057 | 30312 ± 8185* |
| ASVs inferred | 1 229 ± 155 | 1 103 ± 375 | 201 ± 52 | 89 ± 30* |
| Total reads and ASV number | 1 716 653 reads and 7 233 ASVs |  | 1 744 228 reads and 1352 ASVs |  |
| Taxonomy filtering and singletons and doubletons removal | 1 008 592 reads and 5869 bacterial ASVs |  | 560 165 reads and 1065 fungal ASVs |  |

\*Excluding sample 69Mro

For the fungal dataset, we obtained an average of 67 000 raw reads per sample, excluding sample 69Mro. This sample initially contained a far superior number of reads (483 297), which could be due to a volume error added to the ITS pool for this sample (error during quantification) according to Genome Quebec. After the DADA2 filtration steps, we conserved 44.0% and 44.7% of the initial reads for the rhizosphere and root samples respectively (Table 1). Finally, we removed 206 non-fungal ASVs as well as 55 singletons and doubletons resulting in 1065 ASVs and 560 165 reads for 48 samples in our fungal dataset (excluding controls and mock community). In the end, after the filtering process, the 69Mro sample contained 34 665 reads dispatched in 73 ASVs compared to 4739 ± 2215 reads and 68 ± 28 ASVs in the other root samples. Similarly to bacteria, the rarefaction curves obtained for the fungal community indicate that we were able to capture most of the diversity even though more than half of the initial sequences were lost during the filtering process (Supplementary Figure 3).

In terms of negative control, we only found one ASV assigned to *Pezoloma ericae* both present in our dataset and in the rhizosphere extraction blank. This fungus is commonly found in the blueberry root environment. Although this ASV could have originated from a possible contamination from the control and could skew the abundance of this taxon in our dataset, we kept the ASV as it was commonly found across the dataset regardless of the fertilizer treatment. Our pipeline assigned the correct genus for 17 of the 19 species present in the fungal mock community, and 10 of which had a correct species assignment. Although we used the even mock community (equal 18S rRNA gene abundance among strains), Ascomycota taxa were more abundant than Basidiomycota taxa in our sequenced mock.

### Microbial composition

#### Bacteria

The bacterial community was mainly composed of four phyla: Actinobacteriota (1015 ASVs, 36.7% of relative abundance (RA)), Acidobacteriota (720 ASVs, 20.2% RA), Proteobacteria (1199 ASVs, 19.4% RA), and Planctomycetota (1024, 13.3% RA) (Supplementary Figure 4). In these four phyla, one order always stands out: Rhizobiales for Proteobacteria (208 ASVs, 10.0% RA), Frankiales for Actinobacteriota (256 ASVs, 18.1% RA), Acidobacteriales for Acidobacteriota (333 ASVs, 12.9% RA), and Isosphaerales for Planctomycetota (272 ASVs, 9.2% RA). At a finer scale, we observe several taxa with a higher relative abundance than the rest of the community. For instance, the *Roseiarcus* genus (38 ASVs, 2.1% RA) is the most abundant of the Rhizobiales order, as is the case for the *Acidothermus* (210 ASVs, 17.4% RA) and *Mycobacterium* (72 ASVs, 4.8% RA) genera belonging to Actinobacteriota. The Isosphaerales are dominated by the *Aquisphaera* genus (101 ASVs, 6.9 % RA) while the Acidobacteriales order is more evenly distributed with the *Occallatibacter* (70 ASVs, 2.6 % RA), *Acidipila−Silvibacterium* (37 ASVs, 2.6 % RA) and *Granulicella* (65 ASVs, 1.3 % RA) genera. Finally, the *Conexibacter* genus comprised of 142 ASVs, representing 3.1% of the relative abundance, dominates the Solirubrobacterales order while the *Bryobacter* and *Candidatus Solibacter* are two other dominant genera of the Acidobacteriota phylum with 110 AVS, 1.9% RA, and 94 ASVs, 1.6% RA respectively.

#### Fungi

Overall, the fungal community was clearly dominated by the Ascomycota phylum comprised of 794 ASVs summing up to 92.9% of the relative abundance (Supplementary Figure 5). A high proportion of the Ascomycota belongs to the Helotiales order (327 ASVs, 52.8 % RA) followed by the Chaetothyriales order (116 ASVs, 25.5% RA). The Basidiomycota phylum (219 ASVs, 6.7% RA) was dominated by the Agaricales order (84 ASVs, 4.0% RA) while other phyla were represented by fewer ASVs and low relative abundances. At a finer level, a couple of taxa stand out in the Helotiales order: *Pezoloma ericae* (24 ASVs, 16.2% RA), *Oidiodendron maius* (4 ASVs, 9.2% RA), *Phialocephala fortinii* (7 ASVs, 2.8% RA), *Hyaloscypha variabilis* (17 ASVs, 2.8% RA) as well as the *Belonopsis* genus (11 ASVs, 5.0% RA). The Chaetothyriales order is dominated by the Herpotrichiellaceae but the taxonomic resolution is quite poor with 53 out of 71 ASVs with no assigned genus. The Agaricales order is dominated by *Clavaria sphagnicola* (13 ASVs, 2.5% RA).

### Fertilization effect

In the fungal dataset, the PERMANOVA tests indicated a significant effect of fertilization on the fungal communities (except when using the PhILR distance on the productive stage samples) and fertilization explained approximately 10% of the variance in the datasets. Similarly, fertilization was significant for the bacterial datasets regardless of the distance used or the growth stage and it explained 10 to 15% of the variance. For both fungi and bacteria, we found no interactions between plant compartment and fertilization (Table 2 and Supplementary Table 2).

**Table 2.** Effect of plant compartment and fertilization on bacterial and fungal communities in pruning and harvesting years using PERMANOVA test on PhILR distance matrices.

| Factor | df | Pruning |  | Harvesting |  |
| --- | --- | --- | --- | --- | --- |
|  |  | Bacteria | Fungi | Bacteria | Fungi |
| | | $R^2$ • F value (p value) | | | |
| Plant compartments (Pc) | 1 | 0.20 • **5.97 (0.001)** | 0.19 • **4.93 (0.001)** | 0.29 • **9.54 (0.001)** | 0.18 • **4.74 (0.001)** |
| Fertilization (Fe) | 2 | 0.15 • **2.25 (0.003)** | 0.10 • **1.36 (0.049)** | 0.13 • **2.17 (0.015)** | 0.09 • 1.21 (0.112) |
| Pc x Fe | 2 | 0.05 • 0.68 (0.867) | 0.03 • 0.45 (1.000) | 0.04 • 0.67 (0.862) | 0.04 • 0.55 (0.986) |

Note : df, degree of freedom; significant results are in bold

As plant compartments had a stronger on the microbial communities than fertilization, we decided to look at each compartment individually in order to have a clearer interpretation of the fertilization effect. Consequently, there were four combinations between the growth stages and the plant compartments. We used abbreviations: VG (vegetative growth stage) for the pruning year and PD (productive growth stage) for the harvesting year as well as RO and RH for roots and rhizosphere compartments respectively.

#### Fertilization on alpha diversity

In the pruning year, for bacteria, we found a significant difference between the mineral treatment and the control for the Shannon-Weaver index in both compartments, with a higher alpha diversity index for the control (Figure 1A). We observe the similar trend with the Simpson diversity index but the difference between treatments are not significant (data not shown). For fungi, the Shannon-Weaver index of the organic treatment was significantly lower (2.66 ± 0.5) than the control (3.42 ± 0.185) and mineral treatment (3.18 ± 0.185) in the VGxRO combination. No significant differences were observed for the rhizosphere samples (Figure 1B). Finally, in the harvesting year, no significant differences were observed on both alpha diversity indices for fungi or bacteria regardless of the growth stage and compartment combinations analyzed (data not shown).

**Figure 1.**
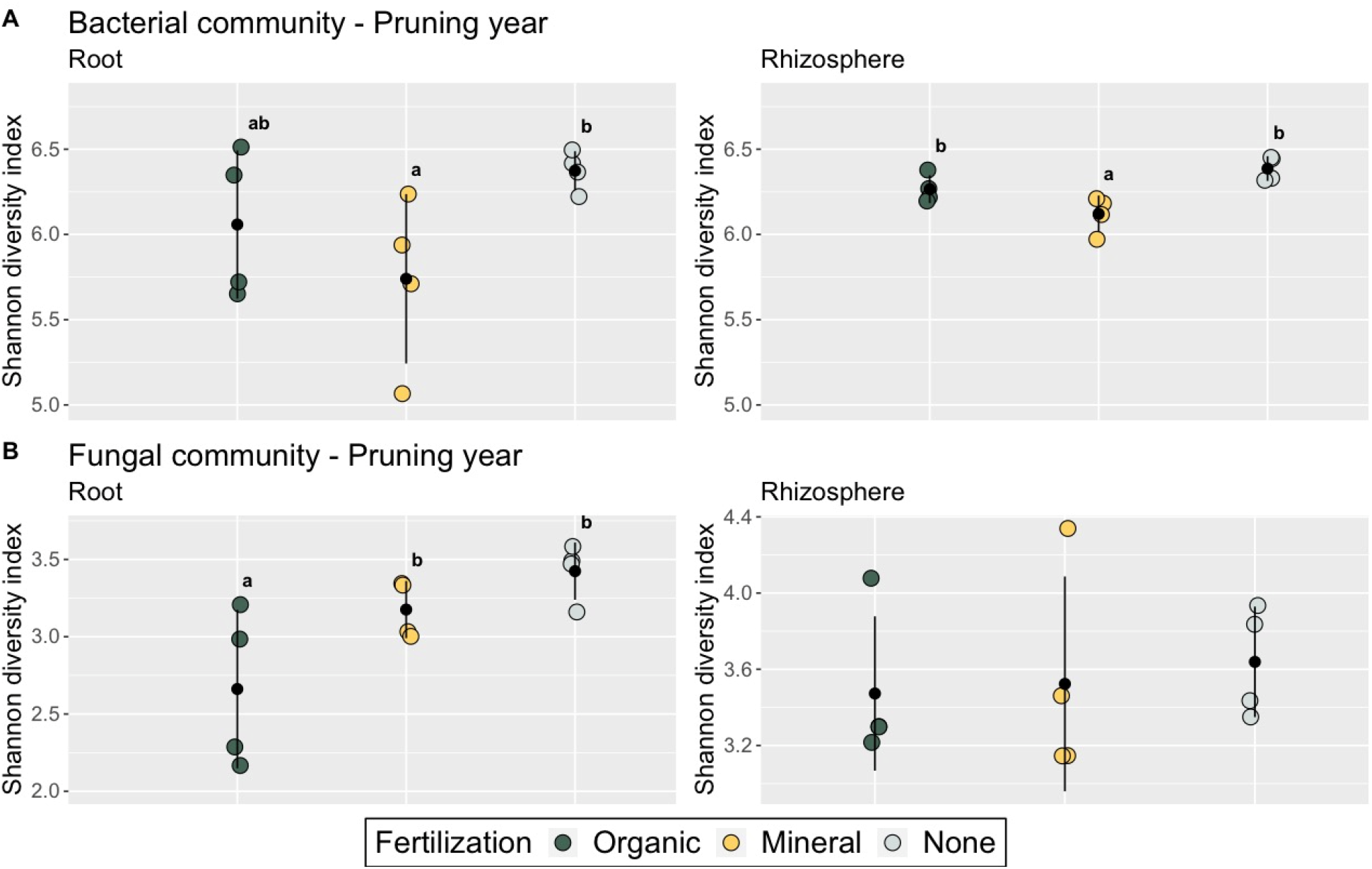
Shannon diversity indices for (A) the bacterial community and (B) the fungal community in the pruning year for each fertilization treatment. Both plant compartments were analyzed with roots in the left column and rhizosphere in the right column. The letters indicate a significant difference between the alpha diversity index found in each plant compartment according to Tukey’s post-hoc test.

#### Fertilization on beta diversity and db-RDA

For the bacterial community, the PERMANOVA test indicated a significant effect of fertilization in both plant compartments in pruning year (VGxRO p = 0.032 for Aitchison, p = 0.025 for PhILR; VGxRH bac p = 0.006 for Aitchison, p = 0.014 for PhILR). The post-hoc tests did not give any significant difference on community composition in the three fertilization treatments, but the difference between the unfertilized and mineral fertilisation were nearly significant for the PhILR distances (VGxRO FDR p= 0.093, VGxRH FDR p = 0.099). Additionally, the ordinations did not show a clear separation of the sample based on their fertilization treatment (Figures 2A and Supplementary Figure 6 A). In the productive growth stage, we found no significant effect of fertilization whether in the root or rhizosphere.

**Figure 2.**
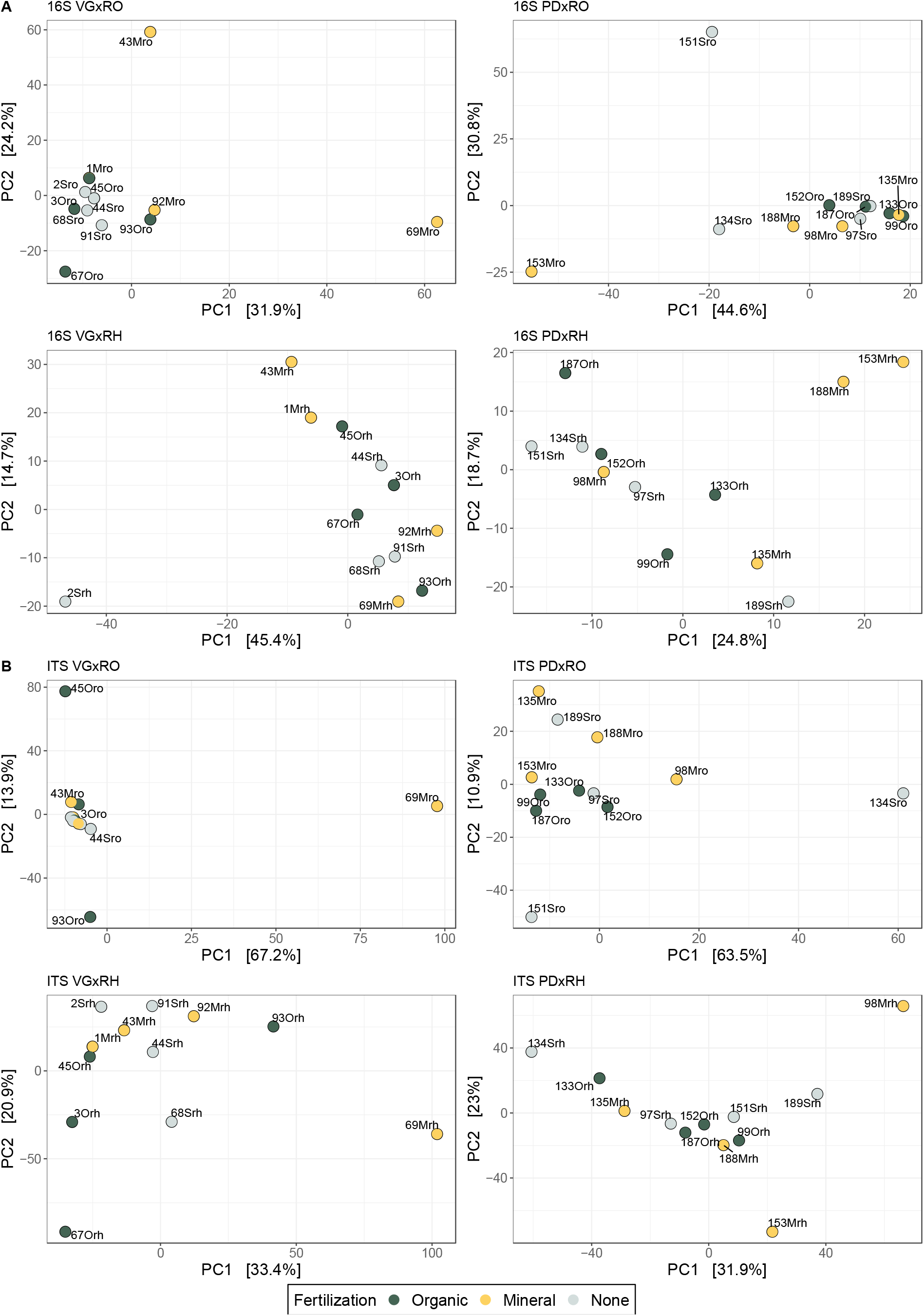
PCA ordinations of (A) the bacterial and (B) fungal communities using the PhILR distance as a representation of beta diversity. All the combinations of plant compartment (RO = roots; RH= rhizosphere) and growth stage (VG = pruning year; PD = harvest year) are presented. Samples are colour-coded according to their fertilizer treatment

Alternatively, the db-RDA constrained by fertilization indicated a poor explanation of the variance in the composition of bacterial communities with very low adjusted R-squared and nonsignificant tests for the productive growth stage. Significant models for the Aitchison distance were found for both compartments in the pruning year (p < 0.05) but the explained variance was low with only 0.4% and 2.3% for the root and rhizosphere bacterial communities respectively.

In the fungal community, we did not find any significant difference in beta diversity, in any of the combinations of plant compartment and growth stage. Similarly to the bacterial community, the samples from each fertilizer treatment overlap and no clear partition is observed (Figure 2 B and Supplementary Figure 6 B). Additionally, similar to what was found for bacteria, the db-RDA analysis showed that the variation in fungal community composition was not redundant with the variance explained by fertilization. The only significant model was found with the Aitchison distance on the pruning year root dataset (p = 0.03) but the adjusted R-squared is extremely low (_adj_R^2^ =0.005).

As a more detailed approach, we used the soil chemistry of the organic layer data as constraining variables in db-RDAs. In the bacterial dataset, using the Aitchison distance, we obtained a significant db-RDA (p = 0.006993) with phosphorus and nitrate as constraining variables for the VGxRO combination though the adjusted R-squared is low (_adj_R^2^ = 0.065). Using the PhILR distance, we find that phosphorus, pH, calcium and nitrate are all significant in the db-RDA and that these variables explain 43.5% of the bacterial community composition (_adj_R^2^ = 0.435, p = 0.001) (Figure 3). Aside from *Mycobacterium cookii* and *Mycobacterium celatum*, the bacteria contributing the most to the community composition of the root samples were not characterized to the species level. Several *Acidothermus spp.* are present on the left side of the graph, in the opposite direction of the pH vector. One *Aquisphaera sp.* seems to play an important contribution in the bacterial composition of root samples which originate from soils with higher P concentration. Similarly, one *Bradyrhizobium sp.* contributes to the bacterial composition of root samples which originate from soils with higher soil Ca and NO_3_ concentrations.

**Figure 3.**
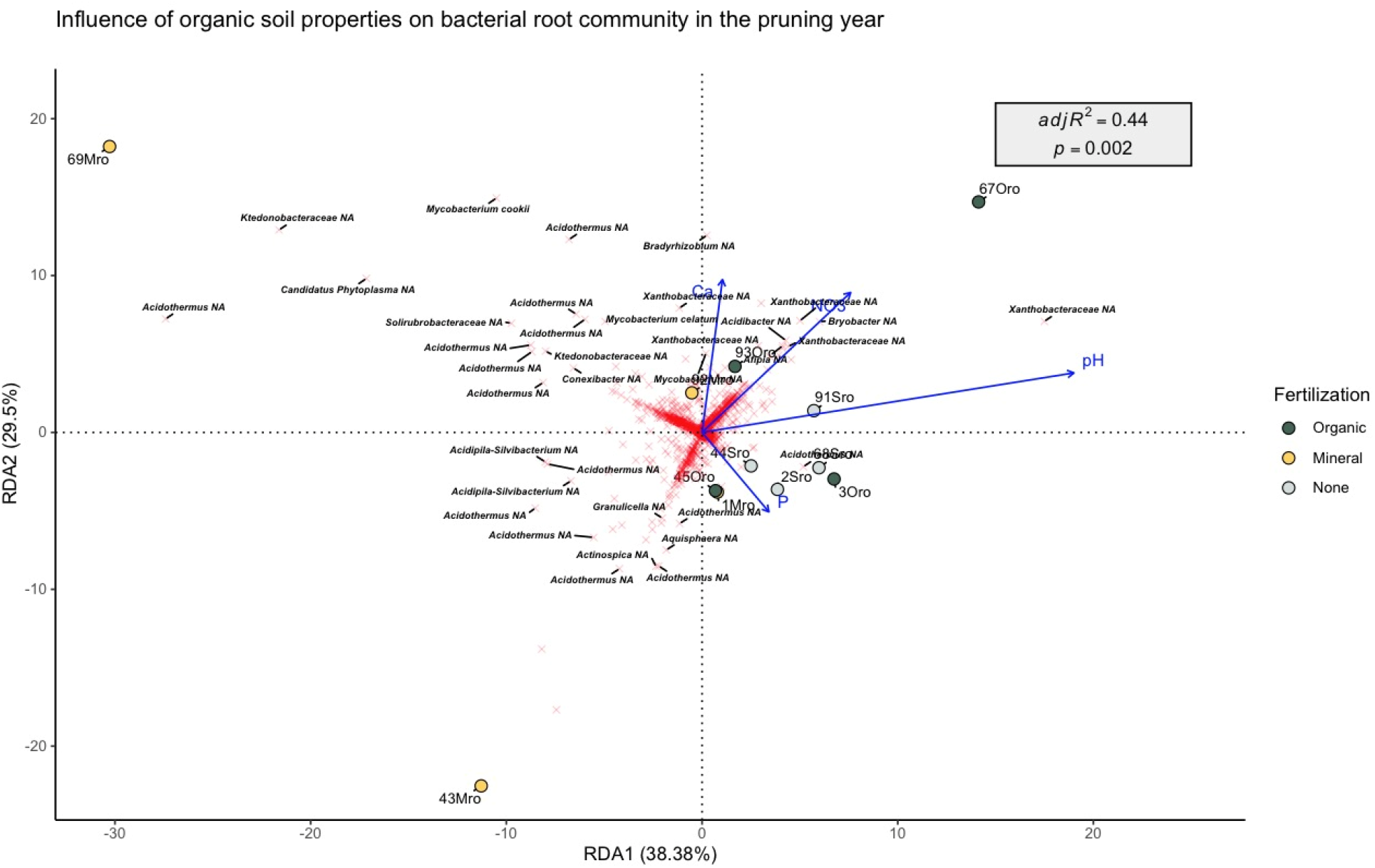
Distance based redundancy analysis (db-RDA) of the bacterial community of the blueberry roots in the pruning year, using the PhILR distance matrix and scaling 1. Blue arrows represent the explanatory variables, red crosses represent the response variables (bacterial ASVs) and the coloured points represent the samples. The perpendicular distance between bacterial ASVs and soil chemistry axes in the plot reflects their correlations, the smaller the distance the stronger the correlation. The response variables (bacterial ASVs agglomerated at species level in this case) were annotated by their taxonomy (up to Family rank) if their scores on either axis were superior to |5| for the sake of clarity.

Additionally, a significant db-RDA was also obtained for the VGxRH combination with phosphorus as the sole constraining variable (p = 0. 03197, _adj_R^2^ = 0. 052) but only for the Aitchison distance (data not shown).

For the fungal datasets, none of the variables were significant when using the Aitchison distance on the four combinations of growth stages and plant compartments. Using the PhILR distance, we found a significant db-RDA for the VGxRH combination, with phosphorus, pH and nitrate as constraining variables (p = 0.006, _adj_R^2^ = 0.31) (Figure 4). Several species stand out, for instance *Pezoloma ericae* plays an important contribution in the fungal community of rhizosphere samples which originate from plots with higher nitrate concentration and pH. On the other end, *Oidiodendron maius* is more present in samples which originate from plots with lower soil pH and phosphorus concentration.

**Figure 4.**
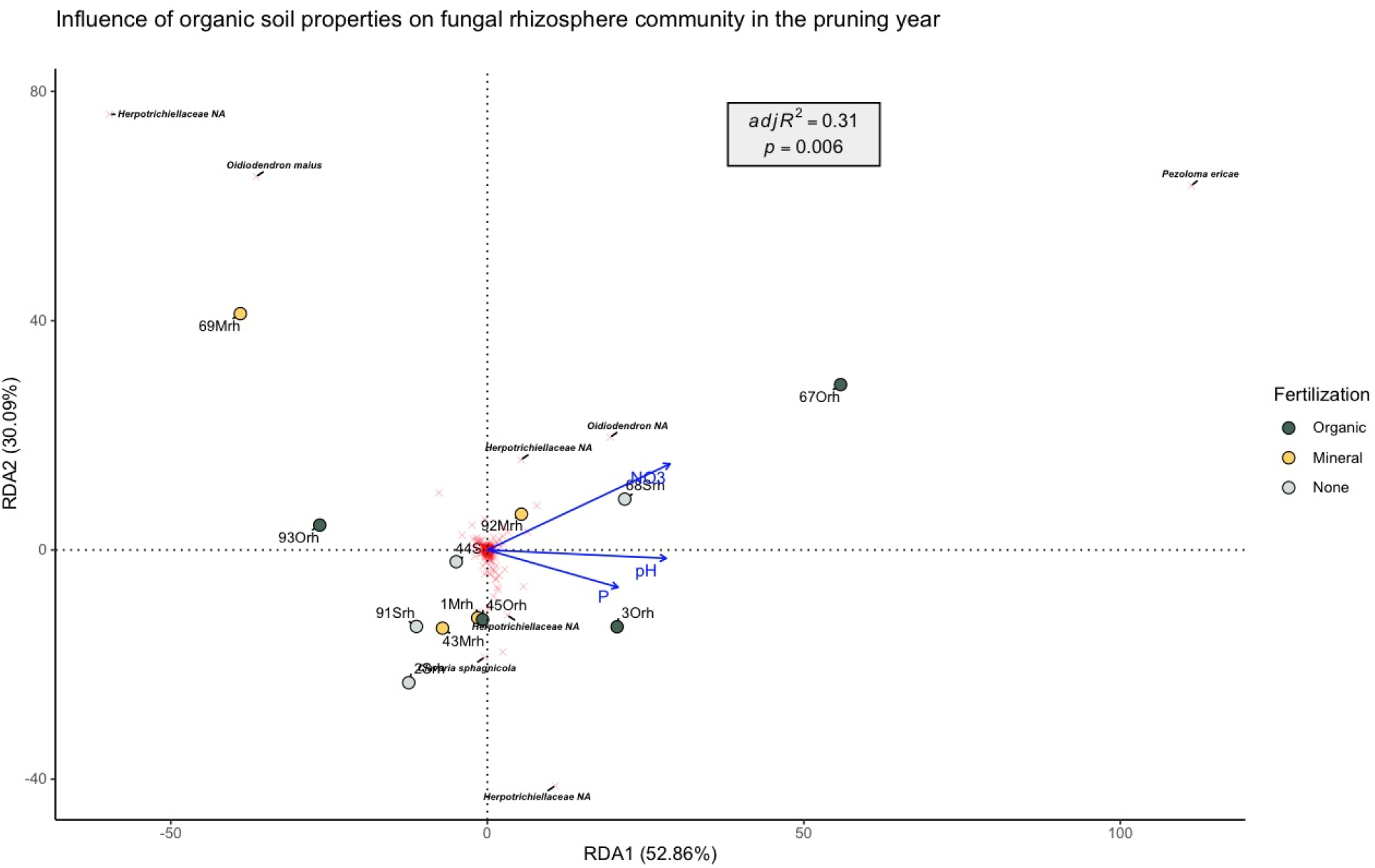
Distance based redundancy analysis (db-RDA) of the fungal community of the blueberry rhizosphere in the pruning year, using the PhILR distance matrix and scaling 1. Blue arrows represent the explanatory variables (soil chemistry), red crosses represent the response variables (fungal ASVs) and the coloured points represent the samples. The perpendicular distance between fungal ASVs and soil chemistry axes in the plot reflects their correlations, the smaller the distance the stronger the correlation. The response variables (fungal ASVs agglomerated at species level in this case) were annotated by their taxonomy (up to Family rank) if their scores on either axis were superior to |10| for the sake of clarity.

#### Fertilization on representative taxa

In order to have a finer approach than the beta diversity analysis which looks at the whole community, we searched for representative taxa in each of the three fertilization treatments in the four combinations of growth stages and plant compartments using two approaches: indicator species and metacoder differential abundance analyses. However, neither analysis found representative taxa of a fertilizer treatment in either the bacterial and fungal communities.

#### Link between agronomic variables and microbiota

As a final analysis, we computed db-RDAs using agronomic variables from both growth stages as constraining variables to test whether the observed variance explained by these variables was redundant with the variance explained by microbial community composition. For fungi, we found no significant results for the harvesting year, and found a weak but significant result for the VGxRO combination using the Aitchison distance (p = 0.04, _adj_R^2^ = 0.07) (Supplementary Figure 7). In this figure, both *Oidiodendron maius* and *Phialocephala glacialis* play an important contribution in samples taken from plots which have lower leaf P and N content contrary to *Mycena sanguinolenta* and *Mollisia cinerea* which are more present in samples taken from plots where samples had an increased leaf P and N. For bacteria, we found a significant result was for the VGxRH combination using the Aitchison distance (p = 0.02, _adj_R^2^ = 0.04) (Supplementary Figure 8). Multiple *Acidothermus spp.* play an important contribution in the bacterial community of rhizosphere samples taken from plots than have higher leaf P and N content.

### Plant compartment effect

#### Alpha and beta diversity

Looking at the bacterial alpha diversity in both roots and rhizosphere, we observed a significant difference for the Simpson index in both the pruning and harvesting growth stages (Kruskal-Wallis test, p < 0.001) with a higher index for the rhizosphere. The mean Shannon-Weavers index is also higher for rhizosphere than for roots but the difference is not significant (Kruskal-Wallis tests, p = 0.069 for the harvesting growth stage, and p = 0.064 for the pruning growth stage) (Figure 5 A). For fungal communities, the only significant difference was for the Shannon Weaver index in the vegetative growth stage, with a lower fungal alpha diversity in the roots compared to the rhizosphere (p = 0.015). There were no significant differences in fungal alpha diversity in the productive growth stage (Figure 5 B).

**Figure 5.**
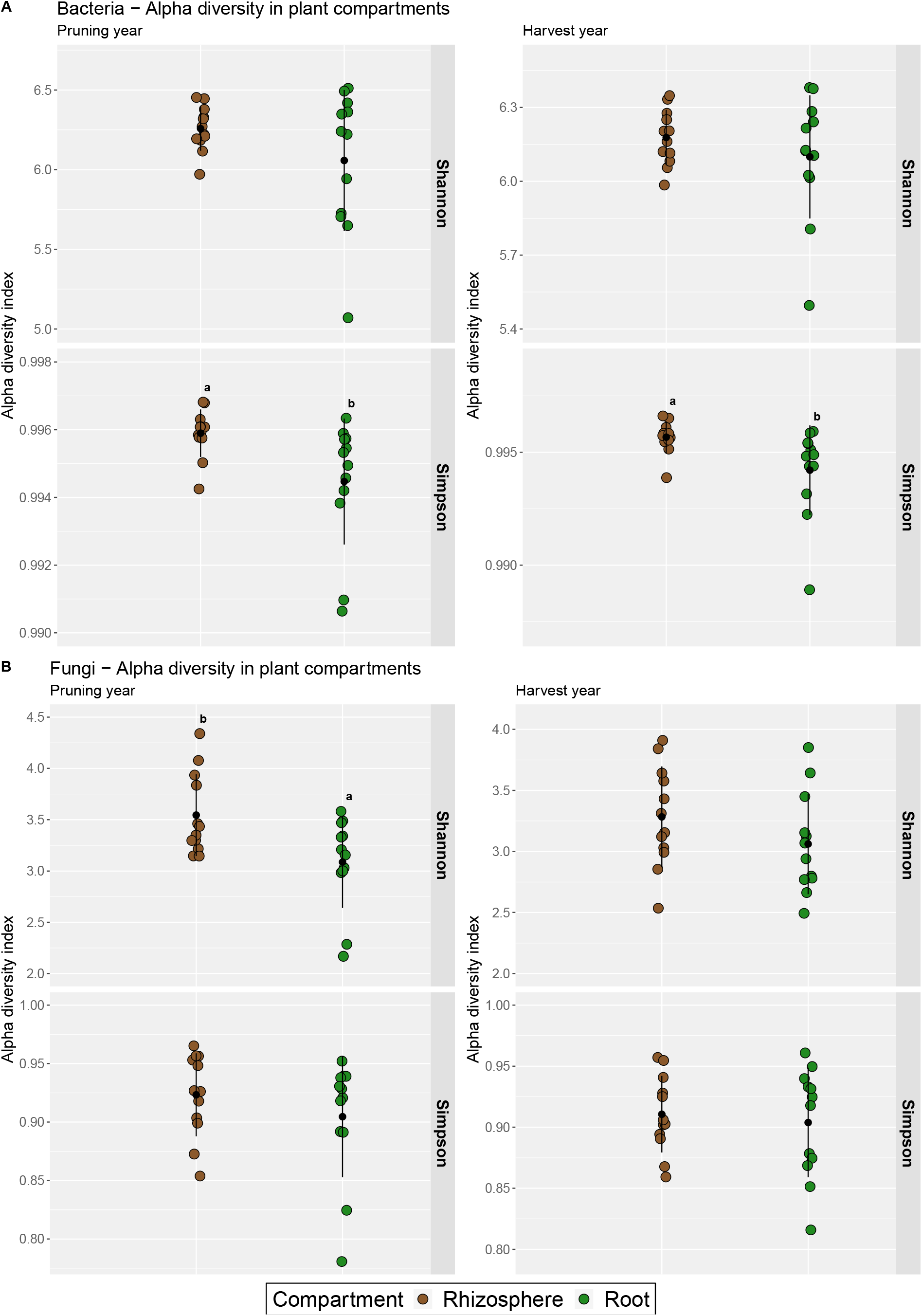
Alpha diversity indices for the bacterial (A) and fungal (B) communities in each plant compartment (brown for rhizosphere, green for roots) for each growth. Both indices Shannon-Weaver and Simpson indices are presented. The letters indicate a significant difference between the alpha diversity index found in each plant compartment.

We obtained significant effects of the plant compartment on the bacterial beta diversity using either the Aitchison and PhILR distances on both the pruning or harvesting subsets (PERMANOVAs p = 0.001) (Figure 6 and Supplementary Figure 9). However, the homogeneity of dispersion assumption was systematically not validated (Betadisper, p <0.05) as shown in the PCA ordinations where the bacterial root community is more dispersed than the rhizosphere (especially when using the PhILR distance) (Figure 6 A). Consequently, the significance of the PERMANOVA tests must be handled with caution, though the visual assessment of the ordinations shows a clear separation between the two plant compartments. Depending on the distance used and on the growth stage, plant compartment explains 10% to 28% of the variance observed. Similarly, we obtained significant effects of the plant compartment on the fungal beta diversity regardless of the growth stage or distance used (PERMANOVAs p = 0.001). For the PhILR ordinations, we observed a higher dispersion of the rhizosphere fungal communities, which was significant for the pruning year (Betadisper, p <0.05) (Figure 6 B). Plant compartment explains 12% to 18% of the variance depending on the growth stage and distance used. The variance explained by each axis of the PCAs are systematically higher in the distance incorporating phylogenetic signals (PhILR).

**Figure 6.**
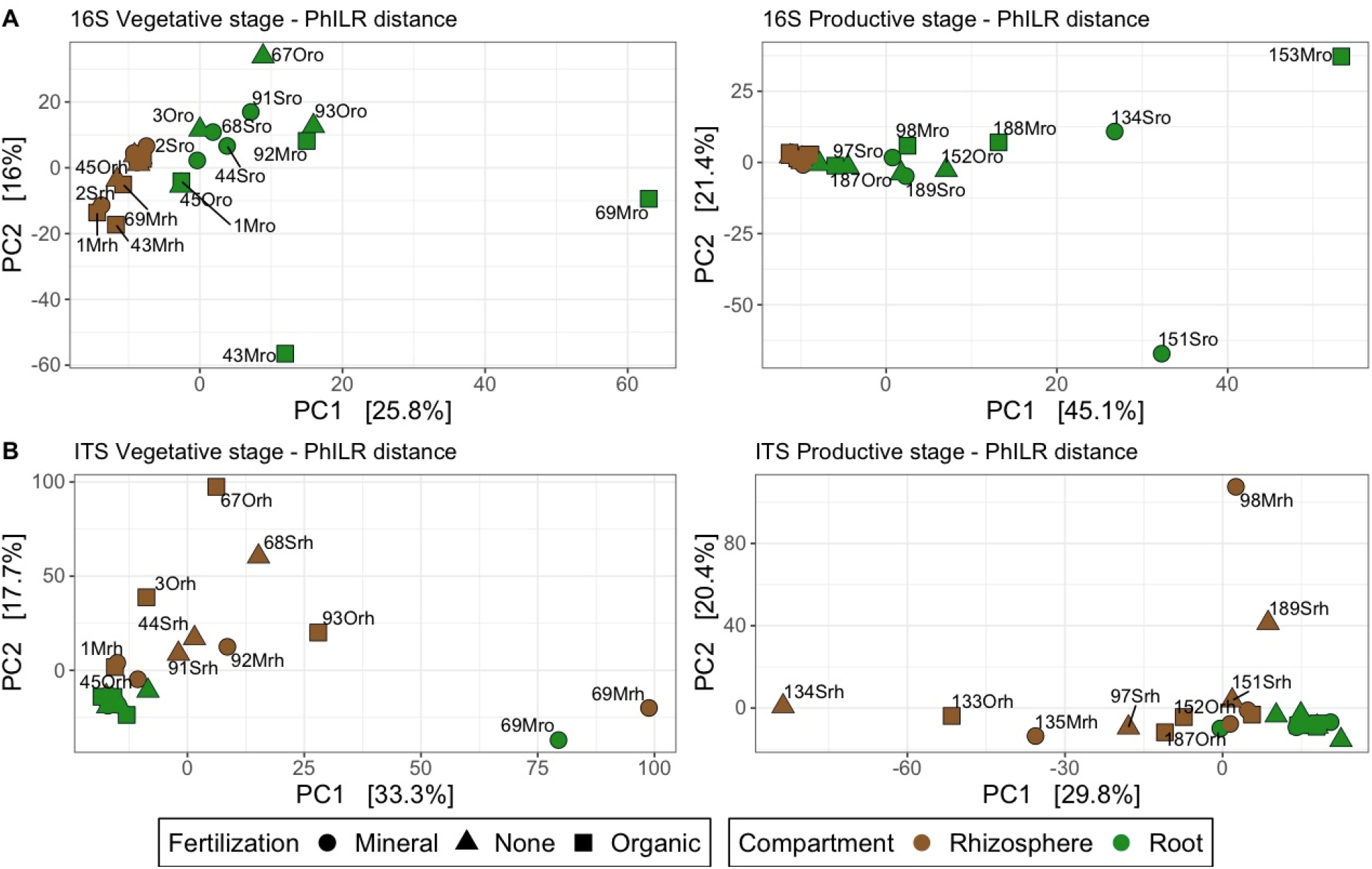
PCA ordinations of (A) the bacterial and (B) fungal communities using the PhILR distance as a representation of beta diversity. Both growth stages are presented (left column: pruning year; right column: harvesting year). The samples are colour-coded according to the plant compartment and shaped according to the fertilization treatment they received.

The differential abundance metacoder plot allows for a finer approach by indicating which taxa are more abundant in a plant compartment. In the pruning year, across the 7 taxonomic levels, we found 56 bacterial taxa that have a significant differential abundance in either plant compartment (Supplementary Figure 10). Overall, the roots are enriched in taxa ranging across multiple lineages including several Gammaproteobacteria while Planctomycetota, and more specifically the Isosphaerales order, are more abundant in the rhizosphere. In the Rhizobiales order, the only significant result concerns the *Methylovirgula* genus which is more abundant in the rhizosphere (log2fold ratio = 1.30, FDR p value = 0.03). In the Acidobacteriales order, the Acidobacteriaceae subgroup 1 is significantly more abundant in the roots (log2fold ratio = 0.85, FDR p value = 0.04). Finally, in the Actinobacteriota phylum, the dominant Frankiales order did not show any significant difference between plant compartments. In the harvesting year, we found a higher proportion of taxa that have a significant differential abundance (107 taxa on 515 in total). The Proteobacteria phylum is significantly more abundant in the roots, with several taxa belonging to Gammaproteobacteria and Alphaproteobacteria having an increased abundance in that plant compartment (Supplementary Figure 11). In the Rhizobiales order, *Bradyrhizobium* spp. are more abundant in the roots (log2fold ratio = 1.72, FDR p value = 0.03) while the *Methylovirgula* genus is more abundant in the rhizosphere (log2fold ratio = 1.21, FDR p value = 0.01). Similarly to the vegetative stage, the Planctomycetota phylum and its Isosphaerales order are significantly more abundant in the rhizosphere.

Looking more closely at this change in fungal community, in the pruning year, we find 80 taxa that have a difference in relative abundance in either plant compartment. Overall, we find numerous taxa belonging to distinct lineages that are exclusive to the rhizosphere compartment (nodes in dark brown in Supplementary Figure 12). In the root compartment, *Belonopsis* sp., *Philocephala fortinii* and *Mycena sp.* are more abundant with log2fold ratios of 7.39, 4.25 and 2.69 respectively (FDR p value < 0.05). In the harvesting year, we obtain 60 individual taxonomic levels with significant changes in relative abundance. The analysis shows that the Helotiales order is significantly more abundant in the roots with a log2fold ratio of 0.66 (FDR p value = 0.023) (Supplementary Figure 13). Similar to what was found in the pruning year, *Belonopsis* sp., *Philocephala fortinii* and the *Mycena* genus were more abundant in the roots with log2fold ratios of 6.74, 3.03 and 3.10 respectively (FDR p value < 0.05). Additionally, *Hyaloscypha variabilis* was also found to be significantly more abundant in the roots (log2fold ratio = 0.45, FDR p value = 0.047). Finally, numerous taxa were found to be exclusive to the rhizosphere, some of which, like *Exophiala xenobiotica* or *Troposporella monospora* were also exclusive to rhizosphere in the pruning year.

## Discussion

### Fertilization has a minimal impact on microbial communities regardless of the growth stage and the compartment

Overall, our results concerning the impact of fertilization on the fungal and bacterial root and rhizosphere communities tend to show a minimal influence in shaping these microbial communities. The only significant differences observed concerned the pruning year samples as expected, because fertilization was applied at the beginning of the pruning year.

For bacteria, we found that the mineral treatment reduces the bacterial Shannon alpha-diversity index in both compartments compared to the control in the vegetative stage. As the Shannon index is more sensitive to rare taxa than the Simpson index, and that there is no significant change for the Simpson index with fertilization, this suggests that mineral fertilization reduces the occurrence of bacterial rare taxa compared to unfertilized plots. As mineral fertilization tend to reduce soil pH, this effect could contribute to explain our result, as pH is one of the principal variables shaping soil microbial communities. Despite this difference in alpha diversity, we did not find any indicative bacterial taxa nor difference in relative abundance for the three fertilization treatments in any of the growth stage and plant compartment combinations. Therefore, we were not able to identify which rare bacterial taxa were depleted due to mineral nutrient addition. We obtained a significant effect of fertilization in both plant compartments for the bacterial beta diversity in the pruning year. However the post-hoc tests did not result in a significant separation of the bacterial communities of the three treatments. Overall, the bacterial community closely resembles to the one described in Morvan *et al*., 2022, as both studies use samples obtained from the same blueberry field and at the same time. Once more, the methanotroph and potentially nitrogen-fixing *Roseiarcus* genus dominates the Rhizobiales order (Kulichevskaya *et al*., 2014), as is the case for the acidophilic *Acidothermus* genus in the Frankiales order, and the Aquispharea genus from the Isosphaerales order known for its carbon-degrading enzymes production (Ivanova *et al*., 2017).

Regarding the fungal community, we observed a significant reduction of the Shannon index for the organic fertilizer treatment in the root compartment during the pruning year while the mineral fertilization Shannon index is not significantly different from the control. We believed that mineral fertilizers would have a stronger impact on the fungal community than the organic fertilizer. The high standard deviation for the organic treatment (two samples with an index value close to 3, more similar to the mineral treatment and control index values and two samples with an index value close to 2.25) sheds some uncertainty on this result. However, organic fertilizer tend to increase soil pH and as the fungal community is clearly dominated by Helotiales, an order containing many acidophilic species, this pH rise could explain the decrease in fungal diversity. Nevertheless, the PERMANOVA tests indicated no significant difference in beta diversity, in any of the combinations of plant compartment and growth stage and we did not find any fungal indicative taxa nor change in relative abundance in any of the three fertilizer treatments. This lack of significance difference in these analyses when comparing the different fertilizer applications show that fertilization does not seem to impact the composition of the fungal community. Finally, similarly to what is observe with the bacterial community, the fungal community is very close to the one observed in Morvan et al., 2022, displaying increased relative abundance of known or putative ericoid mycorrhizal taxa (*Pezoloma ericae*, *Oidiodendron maius*, *Phialocephala fortinii*, *Hyaloscypha variabilis, Clavaria sphagnicola*).

In light of our results, our hypotheses are infirmed as (1) we did not witness a clear composition shift in either microbial communities linked with fertilizer application; (2) the mineral fertilisation did not reduce the occurrence of ErM fungi; and (3) the application of organic fertilization did not reduce the abundance of known ErM fungal species in the roots. Our results are consistent with a previous study, which did not observe any significant change on *Vaccinium* spp. root-associated fungal community composition nor on the assumed ericoid mycorrhizal species under long-term (4 to 12 years) N fertilization (12.5 to 50 kg.ha^-1^) in boreal forests (Ishida and Nordin, 2010). Additionally, testing different rates of mineral N-P-K (from 0 to 60 kg.ha^-1^) in different combinations, Jeliazkova and Percival (2003) found that ericoid mycorrhizal colonization was not impacted by fertilization. Another study, measuring the effect of variable atmospheric nitrogen deposition rates, ranging from 5.27 to 29.65 N kg.ha^-1^.year^-1^ in ombrotrophic bogs containing *Vaccinium oxycoccos*, found that the root fungal diversity (Shannon index) was not impacted by nitrogen deposition rates (Boeraeve *et al*., 2022). The fungal community composition significantly varied with nitrogen deposition rates but this variable did not significantly impact putative ericoid mycorrhizal fungi diversity or composition. In our study, we also obtained sequences matching to known ericoid mycorrhizal species such as *Pezoloma ericae*, *Oidiodendron maius*, *O. chlamydosporicum, Hyaloscypha variablis*, *H. bicolor* and *H. finlandica*. However, we did not find any variation in their relative abundance when comparing the three fertilization treatments. In the same study, the authors also find that increasing nitrogen deposition rates did not significantly change the root bacterial community composition (Boeraeve *et al*., 2022), a pattern similar to what we observe in our experiment. Focusing on the link between fertilization and ericoid mycorrhiza, Kiheri *et al*., (2020) found no significant increase in colonization in *Calluna vulgaris* and *Erica tetralix* (both Ericaceae) in both long-term N deposition and N-P-K fertilization trials (Kiheri *et al*., 2020). Contrasting results to this absence of a fertilization effect have however been found in other studies. In a long-term fertilization experiment (23 to 30 years), Leopold *et al*., (2021) measured a decrease in fungal richness and diversity in the *Vaccinium calycinum* rhizosphere when the soil was supplied with the limiting nutrient (N or P) at a 100 kg.ha^-1^.yr^-1^ dose. In a phosphorus addition trial, Pantigoso *et al*. (2018) measured a shift in bacterial community composition in highbush blueberry rhizosphere when comparing low P (0 and 50 kg.ha^-1^) to high P (101 and 192 kg.ha^-1^) fertilization rates. The authors found an increase in abundance of the Actinobacteria and Bacteroidetes under high P rates while Proteobacteria, Acidobacteria and Verrucomicrobia were more abundant under low P. Although the higher doses of nutrient applied compared to our study could explain these results, in a study on ericoid mycorrhizal richness in several Ericaceae species found in European bogs and heathlands, Van Geel *et al*., (2020) found that increasing nitrogen deposition from 7 to 25 kg.ha^-1^.yr^-1^ and soil phosphorus concentration from 0.02 to 14 mg.kg^-1^ could potentially decrease mycorrhizal richness by 40% and 38%, respectively. Furthermore, when comparing organic and mineral fertilizers applied on *V. corymbosum* at equal doses of N-P-K (60-60-60 kg.ha^-1^), Montalba *et al*. (2010) found that root mycorrhizal colonization was significantly higher with organic fertilizers. The higher mycorrhizal colonization in organic production could be due to the form of nitrogen applied (Sadowsky *et al*., 2012). Unfortunately, we did not quantify mycorrhizal colonization in our study.

The fact that we did not see a strong impact of fertilization on microbial communities could be explained by the limited impact of fertilization on soil chemistry. Indeed, in the pruning year, pH was the only variable that significantly varied between fertilizer treatments (pH for mineral fertilization samples was significantly lower than pH for organic fertilization samples) (Schmitt et al., to be published). Finding a trend of higher pH in the organic treatment and a lower pH in the mineral treatment was to be expected as ammonia tends to acidify the soil (Geisseler and Scow, 2014), while organic fertilizers have already been reported to increase pH (Haynes and Swift, 1985; Caspersen *et al*., 2016). However, the changes in soil chemistry were perhaps too minimal to cause any significant shifts in the fungal and bacterial communities. Another important point to consider is that fertilization occurred in early spring in the pruning year, while soil samples for DNA sequencing were collected approximately 2.5 months after these treatments had occurred. Soil chemistry was measured even later, at the end of September. This lapse of time between fertilization and sampling could be another factor that could explain why we did not find any major changes in the soil ecosystem; the possible initial perturbation of the microbial community caused by fertilization faded with time as the available nutrients were depleted from the organic soil layer. In the harvesting year, the time between fertilization and sampling are even more temporarily apart. A time-series experiment with regular sampling during the summer could show the dynamic of soil chemistry, and microbial community changes caused by fertilization. This type of study would allow to show if the impact of fertilization on the microbial communities is short-lived (<2.5 months) or if the microbial community is resistant to these changes of conditions. However, some of the results obtained with the db-RDAs constrained by soil chemistry make sense such as the bacterial genus *Acidothermus* being more abundant in samples originating from plots with lower pH (Figure 3). This genus has a single species described to date, *A. cellulolyticus,* isolated form the Yellowstone hot springs, and whose genome was sequenced (Barabote *et al*., 2009). As the name implies, this bacteria is both thermophilic and acidophilic. The thermophilic aspect may be surprising in a wild blueberry soil but temperature records show that soil temperature in this blueberry field often reaches 40°C, with a max temperature of 53.07°C in the summer 2021. *A. cellulolyticus* was also characterised as producing several carbohydrate active enzymes, including chitinase (Barabote *et al*., 2009). Despite the low amount of variation explained in the db-RDA constrained by leaf chemistry, the fact that *Acidothermus* spp. play an important contribution of the bacterial community in samples with higher leaf P and N content, could be linked to this hydrolytic activity (Supplementary Figure 8).

### Bacterial and fungal diversity and community composition change with plant compartments

Regarding bacteria, we found a significant difference in the Simpson diversity index between rhizosphere and roots in the samples from both growth stages. However, no significant difference was found for the Shannon diversity index. These results are in line with numerous studies which find a higher diversity in the rhizosphere compared to the roots as the plant selection pressure is increased in its organs (Reinhold-Hurek *et al*., 2015; van der Heijden *et al*., 2015).

We also obtained significant community shifts between rhizosphere and roots for both the Aitchison and PhILR distances and a clear separation of both communities in the ordinations using these distances. The bacterial root samples tend to contain more phylogenetically distant taxa than the rhizosphere as samples are more dispersed in the ordinations plots based on the PhILR distance. Finally, looking at the most abundant phyla, the Metacoder plots indicate a higher relative abundance of Planctomycetota (P) – Isosphaerales (O) in the rhizosphere while several Proteobacteria are enriched in the roots. Species belonging to the Isosphaerales order contain numerous carbohydrate-active enzymes suggesting their ability to use a wide range of natural carbohydrates (Ivanova *et al*., 2017). Several of these Isosphaerales including the Aquispharea genus were also found in abundance in the wild blueberry rhizosphere in a previous study (Morvan *et al*., 2022). As wild blueberry roots are mainly found in the soil organic layer, finding an increased abundance in hydrolytic bacteria in the rhizosphere is coherent. *Bradyrhizobium* spp. were more abundant in the blueberry roots than in the rhizosphere in the productive year. This genus is widely known for its nitrogen fixation capacity in legume nodules. Nevertheless, this genus is also found as an endophyte in non-legumes roots and may contribute to N2 fixation (Yoneyama *et al*., 2017; Rosenblueth *et al*., 2018; Hara *et al*., 2019) but can also show an absence of nitrogen contribution (Schneijderberg *et al*., 2018). Hence, an in-depth characterization of the *Bradyrhizobium* spp. present in the blueberry roots is required to elucidate their nitrogen fixation ability. The *Methylovirgula* genus more abundant in the blueberry rhizosphere than in the roots, was isolated in 2009 from beechwood blocks at the surface of an acidic (pH 3.3–3.6) forest soil and the two isolates were also able to fix dinitrogen (Vorob’ev *et al*., 2009).

For the fungal community, alpha diversity was only significantly higher in the rhizosphere of the vegetative stage samples with the Shannon index, suggesting that it was enriched in low abundance taxa. Fungal community composition was also significantly different when comparing plant compartments as it can be seen on the ordinations. Contrary to bacteria, the fungal root samples tend to have communities that are more phylogenetically close to one another when the rhizosphere communities are more phylogenetically different. In both growth stages, many fungal taxa are exclusive to the rhizosphere including known ericoid mycorrhizal fungi such as *Oidiodendron chlamydosporicum* and *Hyaloscypha finlandica*. However the dark septate endophyte *Phialocephala fortinii* as well as the *Mycena* genus are consistently more abundant in the roots. *P. fortinii* commonly colonizes Ericaceae roots and could have a beneficial effect on their fitness (Vohník *et al*., 2005; Newsham, 2011; Lukešová *et al*., 2015). As for the Mycena genus, Grelet *et al*., (2017) have proposed that this genus, previously characterized as a generalist saprotroph, could be transitioning between a full saprotrophic and a symbiosis lifestyle. The authors obtained a comparable increase of growth in seedlings of *Vaccinium corymbosum* inoculated with either *Mycena galopus* or *Pezoloma ericae*, and observed peg-like structure in the root cells inoculated with *M. galopus*. However, as we did not look at colonization, we cannot confirm if we had a similar structure in the *Vaccinium angustifolium* roots that served to extract DNA. Another *Mycena* species, *M. sanguinolenta*, was also found to play an important contribution to the fungal community of samples with a higher P and N leaf content, along with *Mollisia cinerea* (Supplementary Figure 7). *Mollisia* is an understudied fungal group that contain a range of lifestyles from saprotrophs to endophytes, similar the phylogenetically close Phialocephala genus (Tanney and Seifert, 2020).

## Conclusion

The results of our study indicate a minimal impact of fertilization on both the fungal and bacterial communities of the rhizosphere and roots of *Vaccinium angustifolium*. Fertilization is clearly worthwhile for producers as yields strongly increase when the mineral and organic fertilizers are used. Our experiment did not measure drastic changes in soil chemistry in the soil organic layer, as pH was the only variable which significantly varied in the pruning year when comparing the two fertilizer treatments (significantly lower pH with the mineral fertilizer compared to the organic fertilizer). As fertilization is applied at a much lower rate compared to other crops, this could explain why the shifts in microbial community observed in other settings did not occur in this experiment. A long-term impact of fertilization would be interesting to measure, as the plots sampled were relatively recent in terms of agricultural management. Repeated nutrient addition could progressively modify soil chemistry which in turn could impact the microbial communities in the root environment of cultivated wild blueberries. Furthermore, although we found no relations between yield and the microbial communities, future studies based on functional omics approaches would allow to have a more detailed understanding of the contribution of nitrogen fixing bacteria, ericoid mycorrhizal fungi and dark septate endophytes in the wild blueberry nutrition.

## Author contribution

MP and JL conceived and designed the agronomic part of the study while MH and SM conceived and designed the microbial part of the study. SM contributed to data acquisition, performed the microbial community analysis experiments, analyzed the data and wrote the draft of the manuscript. All authors read and approved the submitted version of the article.

## Acknowledgments

This study received funding from the following source: the Natural Sciences and Engineering Research Council of Canada (NSERC) Discovery grant (Grant RGPIN-2018-04178) to MH; the Syndicat des Producteurs de Bleuets du Québec (SPBQ) and the Natural Sciences and Engineering Research Council of Canada (NSERC) (Grant RDCPJ-503182-16) to MP.

The authors declare that the research was conducted in the absence of any commercial or financial relationships that could be construed as a potential conflict of interest.

The authors would like to thank Geneviève Telmosse for her help on data acquisition; Denis Bourgault for soil analysis; Stéphane Daigle for his help on mixed models; and Andrew Blakney for his support. The authors also thank the Corporation d’Aménagement Forêt Normandin (CAFN) for providing access to their sites and infrastructure.

## Supplementary Information

**Supplementary Figure 1.**
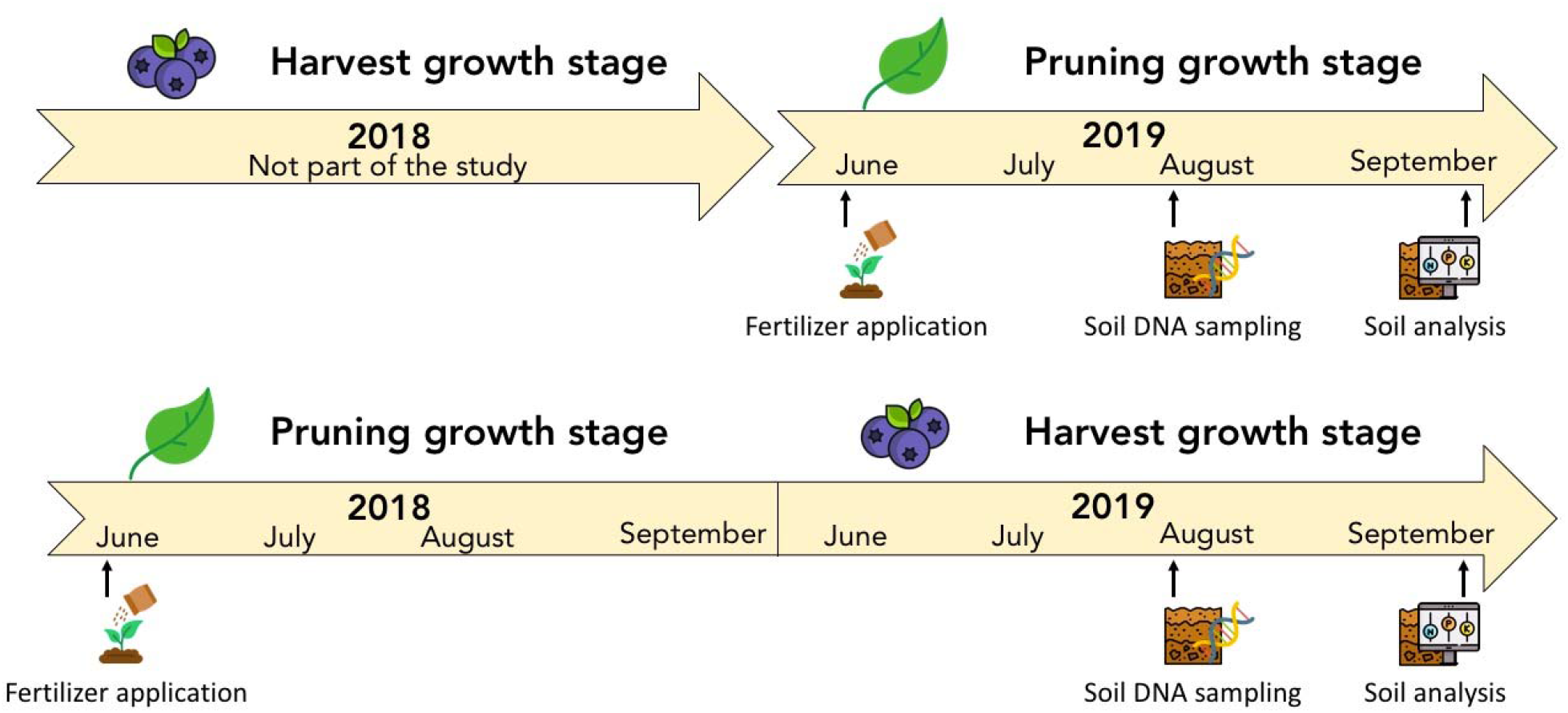
Graphical abstract

**Supplementary Figure 2.**
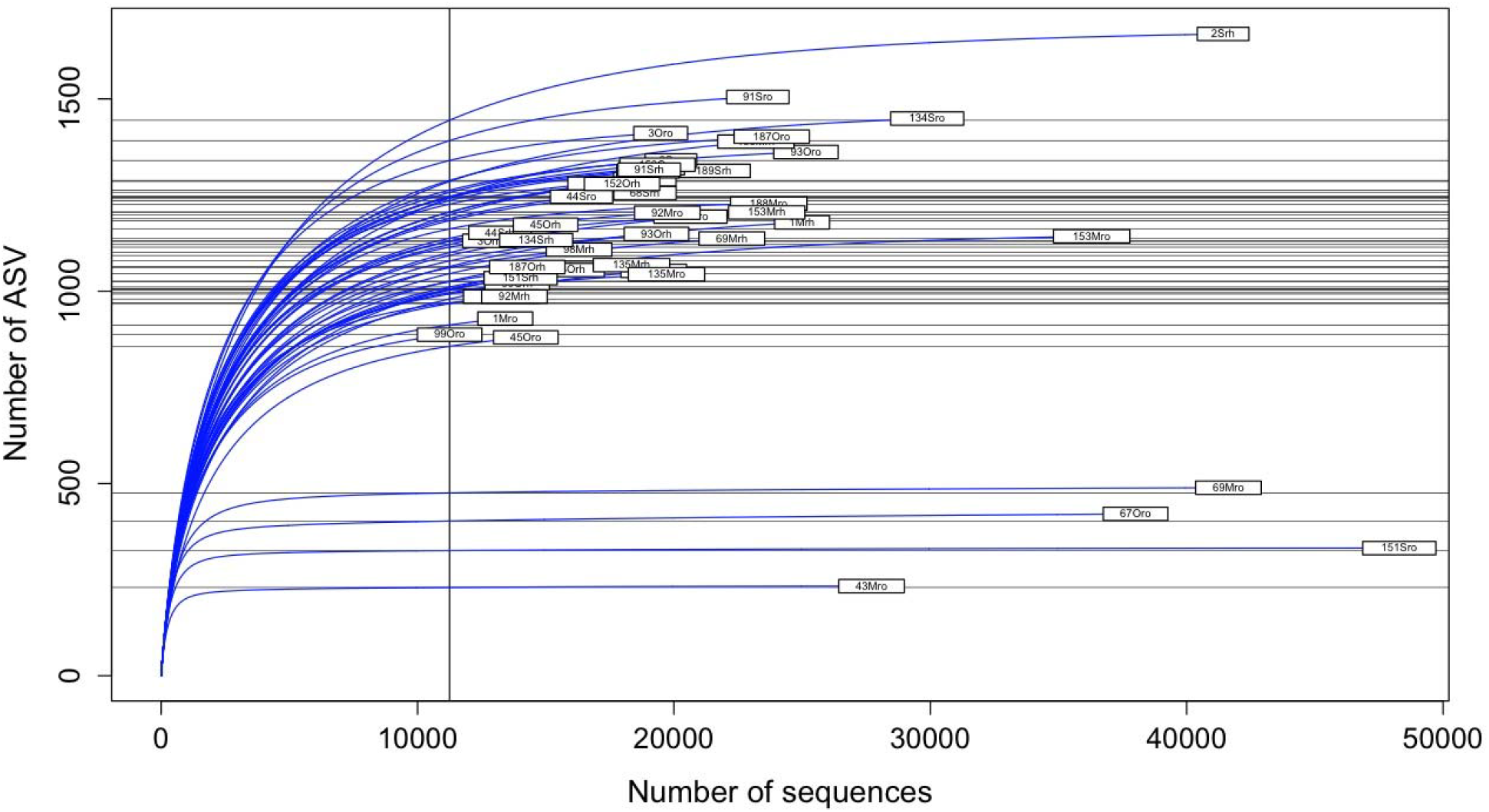
Rarefaction curves of the bacterial community after the filtering pipeline.

**Supplementary Figure 3.**
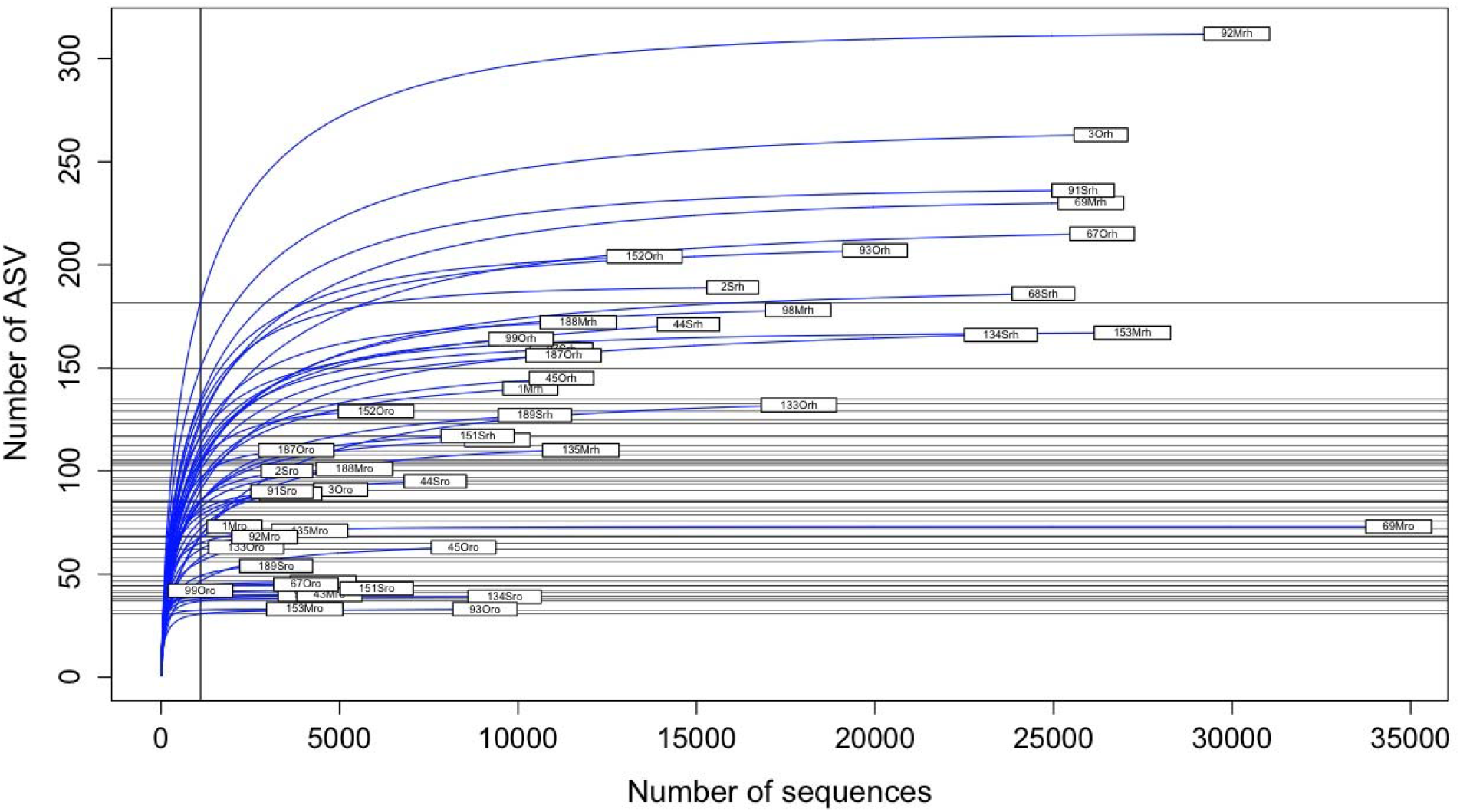
Rarefaction curves of the fungal community after the filtering pipeline.

**Supplementary Figure 4.**
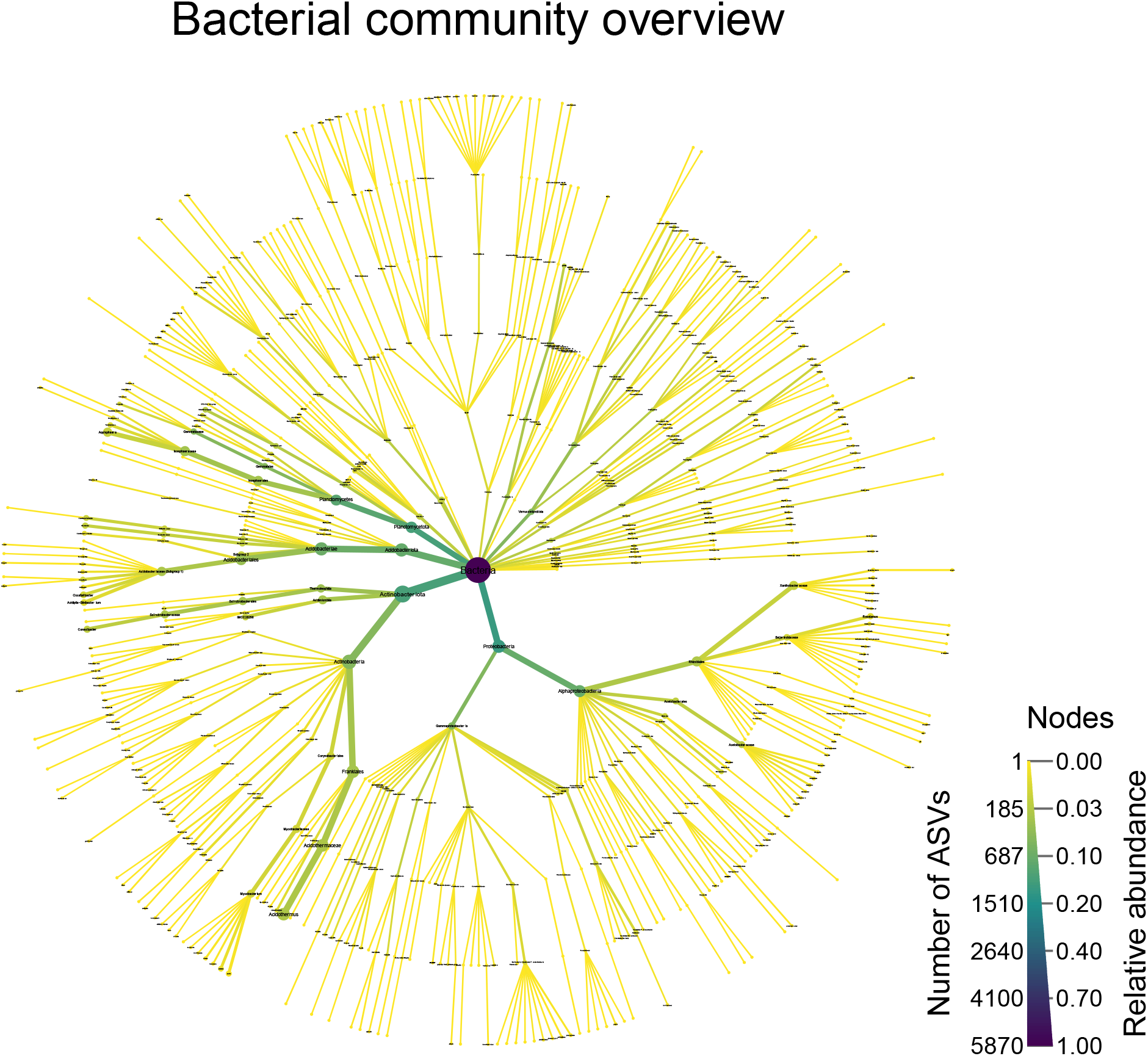
Taxonomic overview of the bacterial community. The figure plots ASVs which were assigned to a taxonomic level (ASVs not assigned at a particular taxonomic level (NAs) are not shown). The colour of the edges and nodes indicates the number of ASVs found at a given taxonomic level, with darker colour indicating more ASVs and lighter colours indicating fewer ASVs. The size of the edges and nodes indicate the relative abundance of the ASVs assigned to a particular taxonomic level with wider edges/nodes indicating a high relative abundance and narrower edges/nodes indicating a low relative abundance.

**Supplementary Figure 5.**
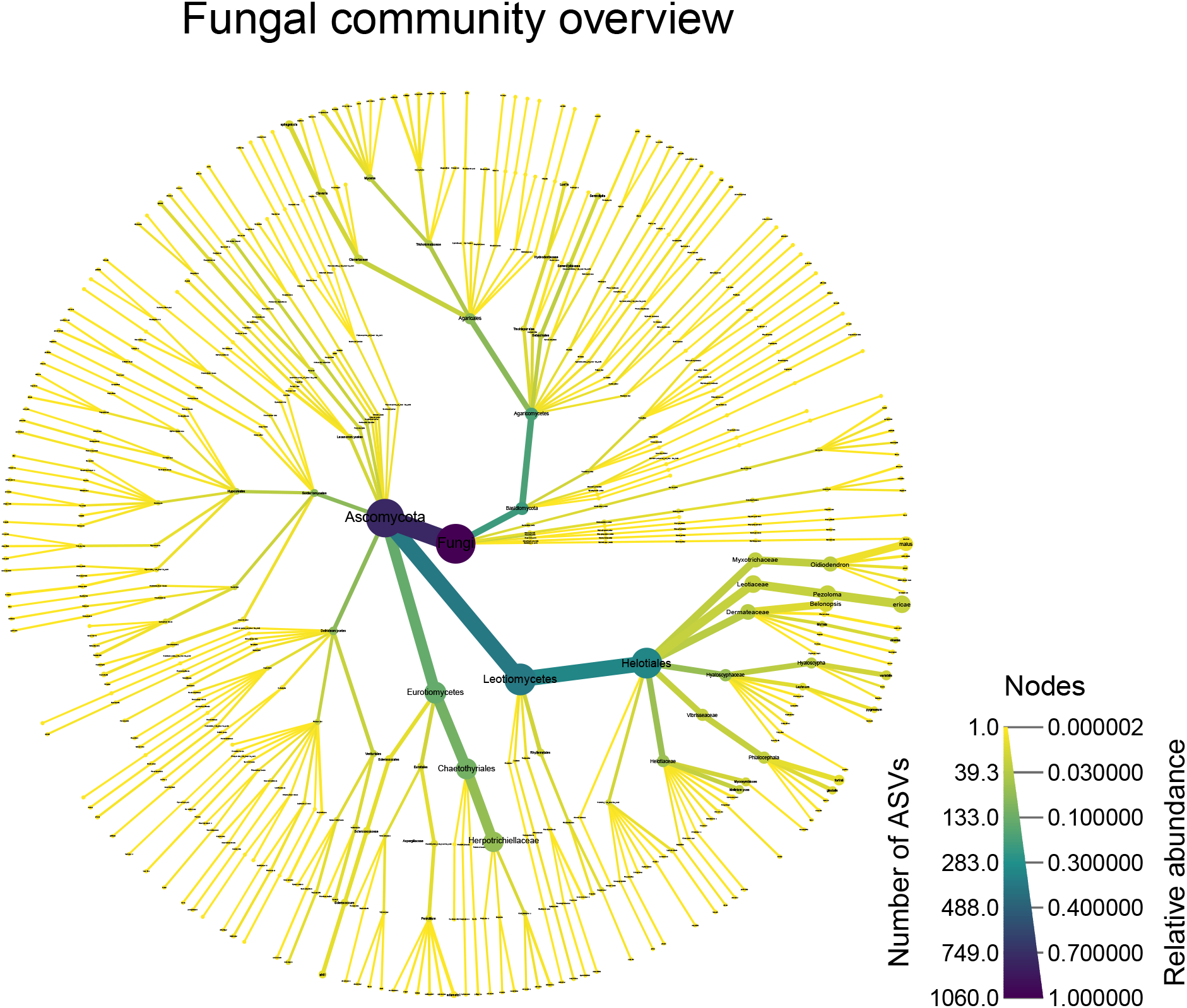
Taxonomic overview of the fungal community. The figure plots ASVs which were assigned to a taxonomic level (ASVs not assigned at a particular taxonomic level (NAs) are not shown). The colour of the edges and nodes indicates the number of ASVs found at a given taxonomic level, with darker colour indicating more ASVs and lighter colours indicating fewer ASVs. The size of the edges and nodes indicate the relative abundance of the ASVs assigned to a particular taxonomic level with wider edges/nodes indicating a high relative abundance and narrower edges/nodes indicating a low relative abundance.

**Supplementary Figure 6.**
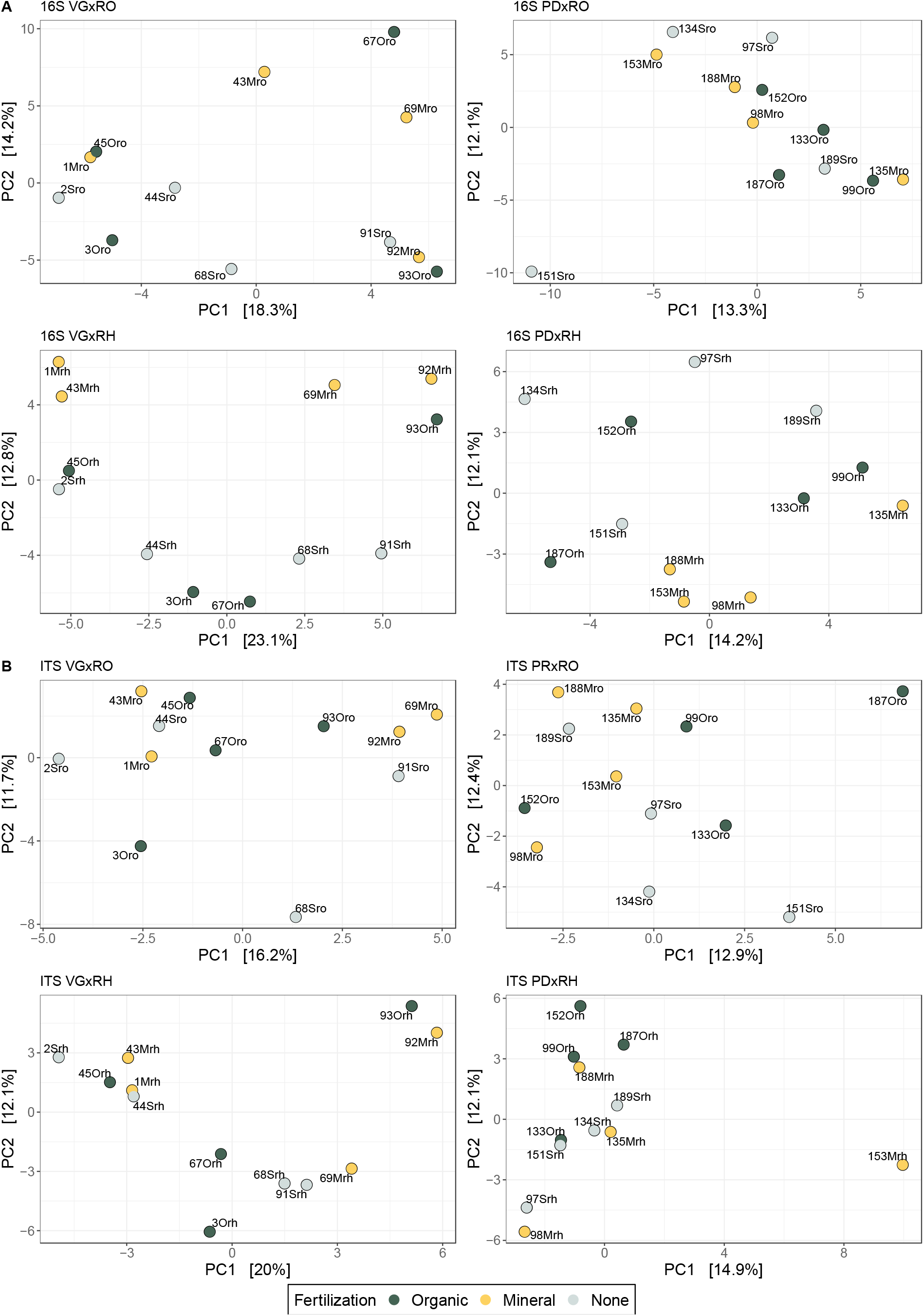
PCA ordinations of (A) the bacterial and (B) fungal communities using the Aitchison distance as a representation of beta diversity. All the combinations of plant compartment (RO = roots; RH= rhizosphere) and growth stage (VG = pruning year; PD = harvest year) are presented. Samples are colour-coded according to their fertilizer treatment.

**Supplementary Figure 7.**
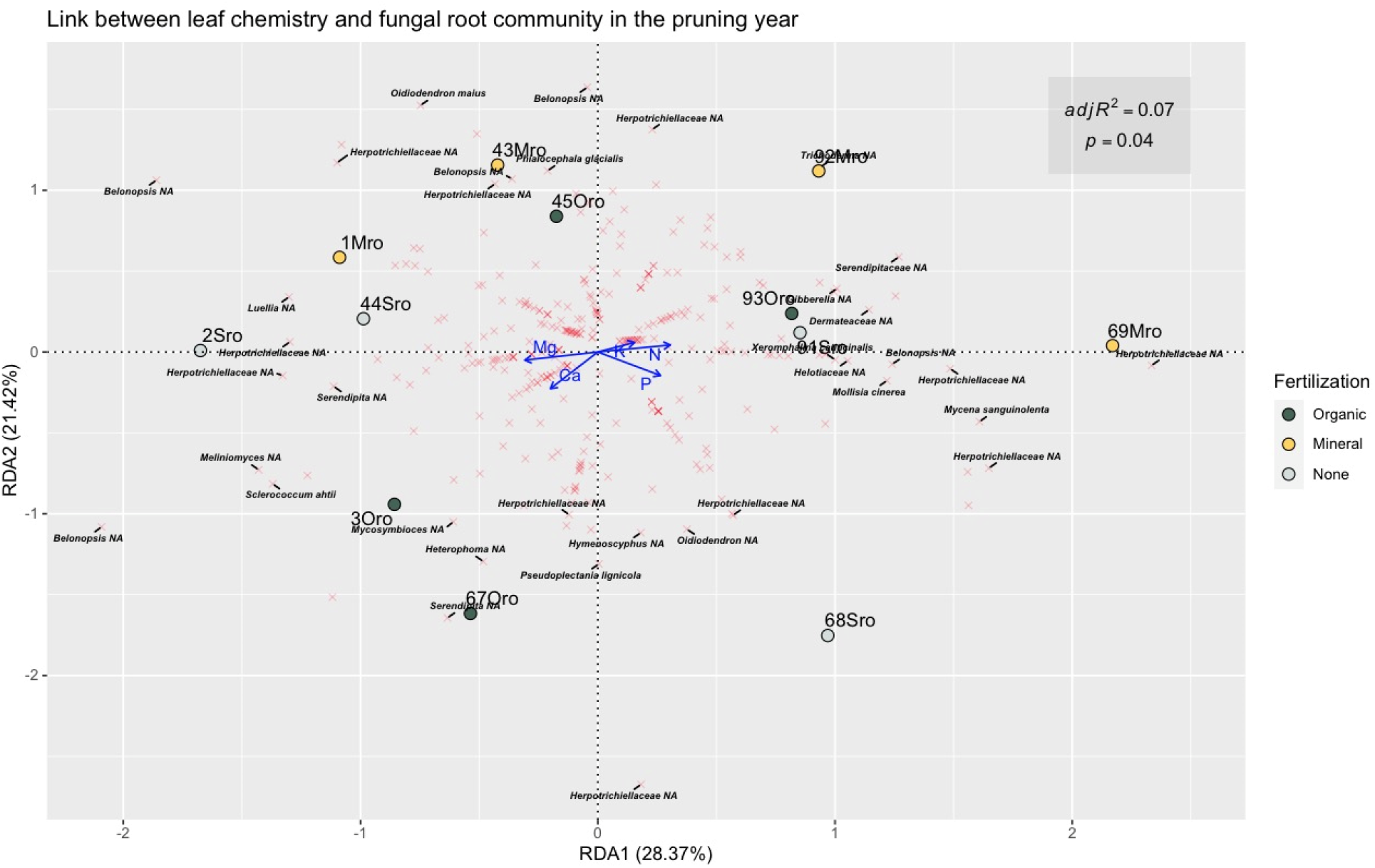
Distance based redundancy analysis (db-RDA) of the fungal community of the blueberry roots in the pruning year, using the PhILR distance matrix and scaling 1. Blue arrows represent the explanatory variables (leaf chemistry), red crosses represent the response variables (bacterial ASVs) and the coloured points represent the samples. The perpendicular distance between bacterial ASVs and soil chemistry axes in the plot reflects their correlations, the smaller the distance the stronger the correlation. The response variables (fungal ASVs agglomerated at species level in this case) were annotated by their taxonomy (up to Family rank) if their scores on either axis were superior to |1| for the sake of clarity.

**Supplementary Figure 8.**
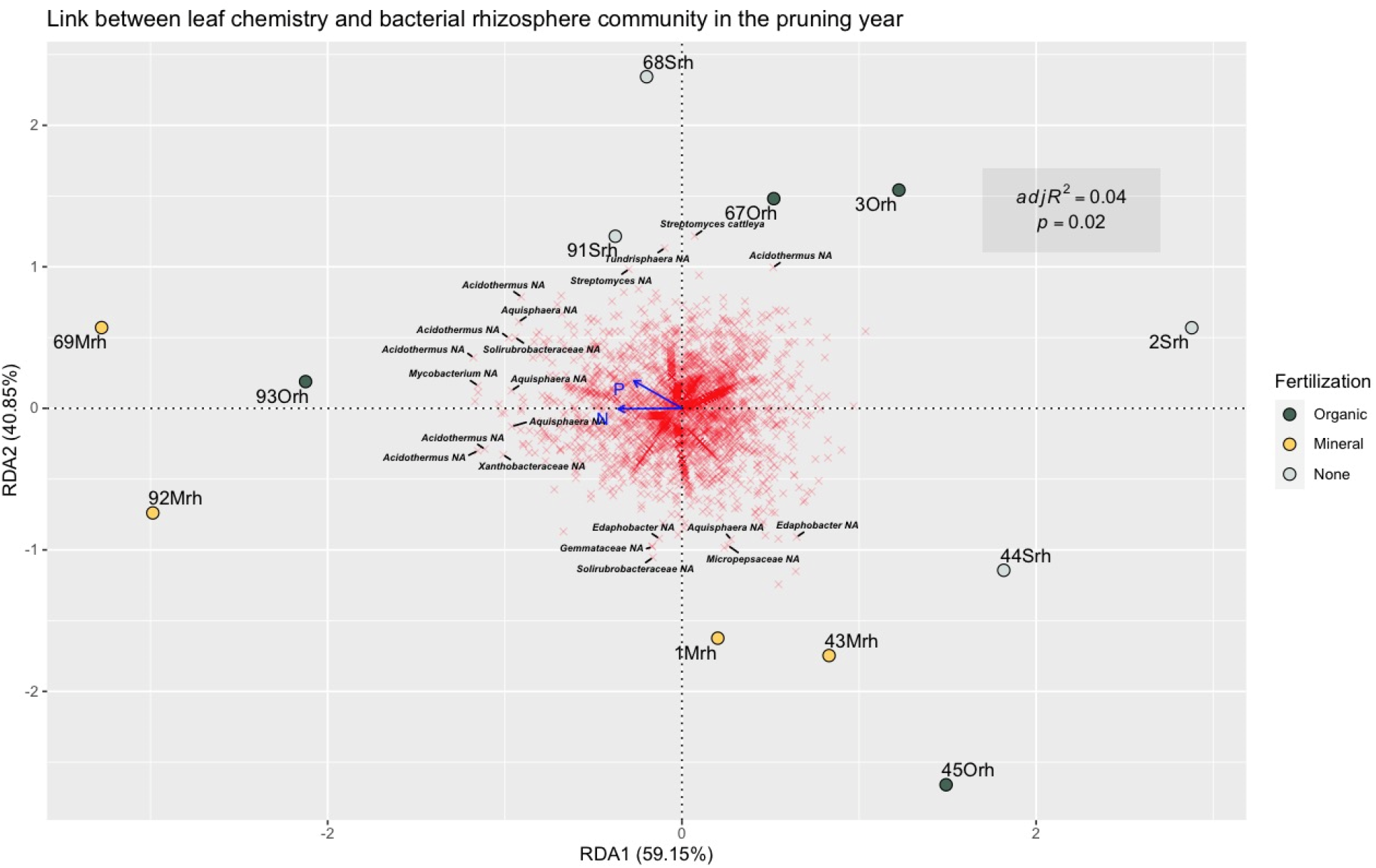
Distance based redundancy analysis (db-RDA) of the bacterial community of the blueberry rhizophere in the pruning year, using the PhILR distance matrix and scaling 1. Blue arrows represent the explanatory variables (leaf chemistry), red crosses represent the response variables (bacterial ASVs) and the coloured points represent the samples. The perpendicular distance between bacterial ASVs and soil chemistry axes in the plot reflects their correlations, the smaller the distance the stronger the correlation. The response variables (bacterial ASVs agglomerated at species level in this case) were annotated by their taxonomy (up to Family rank) if their scores on either axis were superior to |1| for the sake of clarity.

**Supplementary Figure 9.**
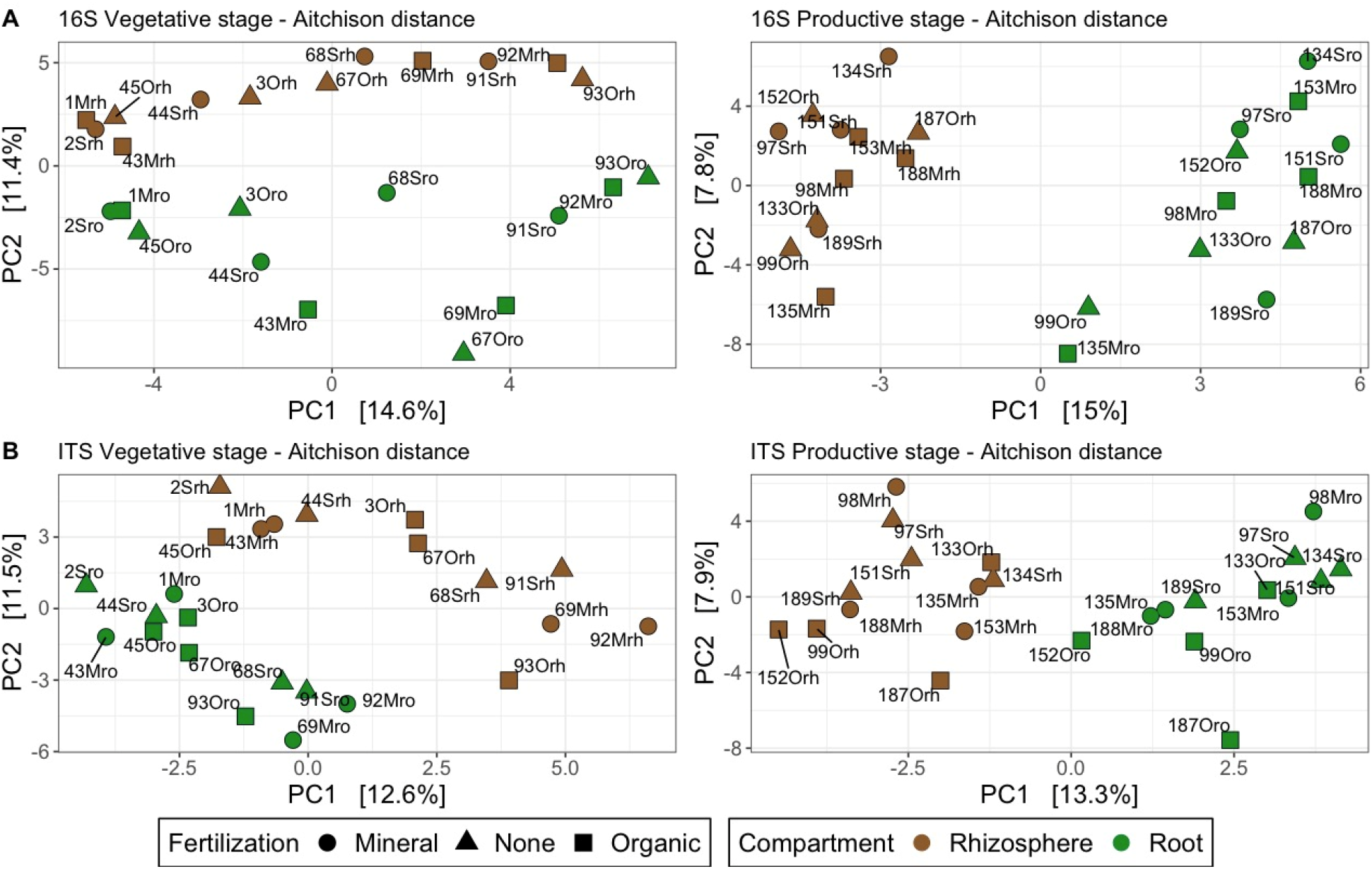
PCA ordinations of (A) the bacterial and (B) fungal communities using the PhILR distance as a representation of beta diversity. Both growth stages are presented (left column: pruning year; right column: harvesting year). The samples are colour-coded according to the plant compartment and shaped according to the fertilization treatment they received.

**Supplementary Figure 10.**
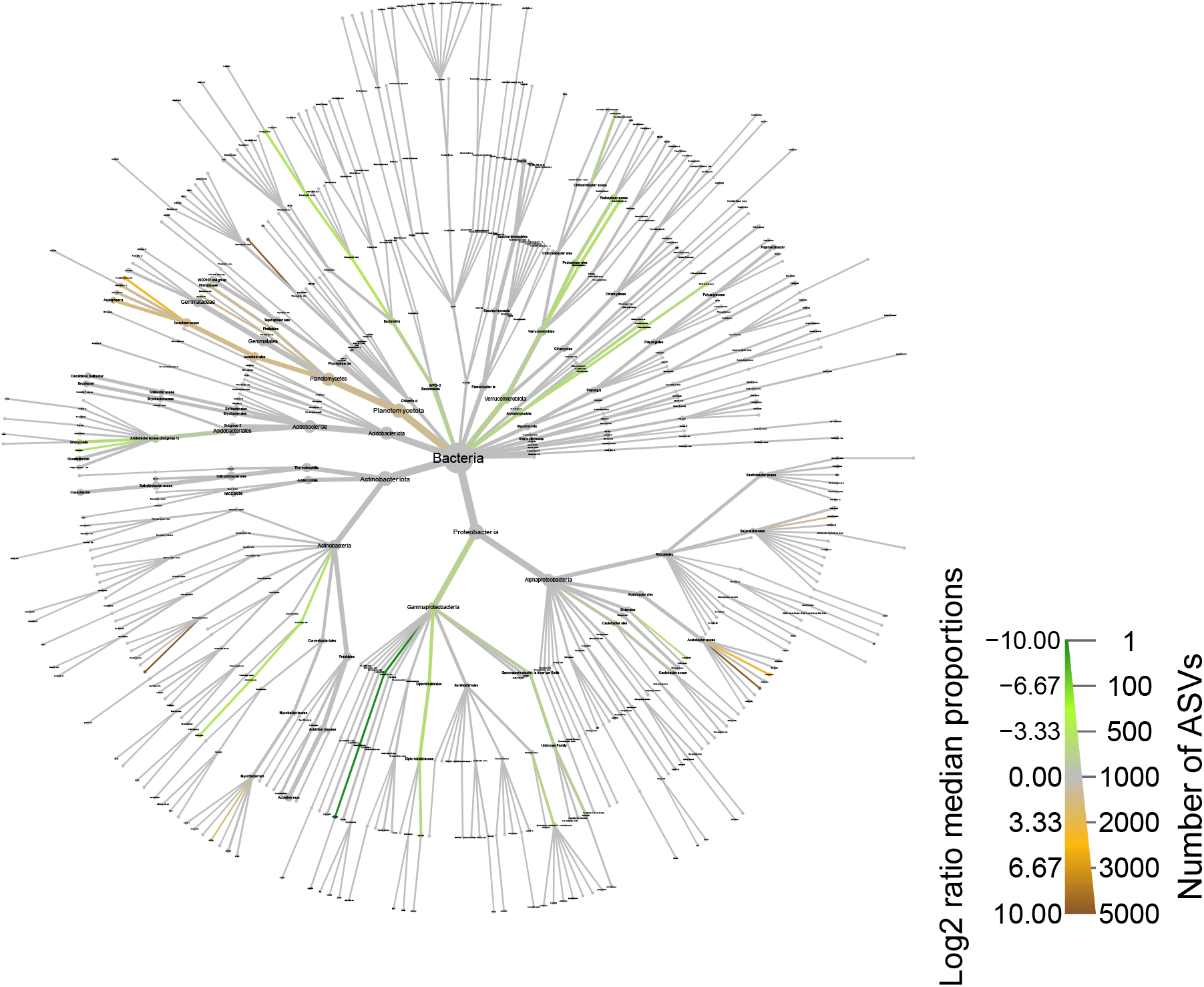
Differential abundance in the bacterial community for the pruning year comparing the root and rhizosphere. The coloured ASVs have a significant difference in relative abundance in either plant compartment (green for roots, brown for rhizosphere). A log2ratio of 10 (dark brown) or −10 (dark green) indicates that the taxon was exclusively present in the given plant compartment. The size of the edges and nodes indicate the number of ASVs assigned to a particular taxonomic level.

**Supplementary Figure 11.**
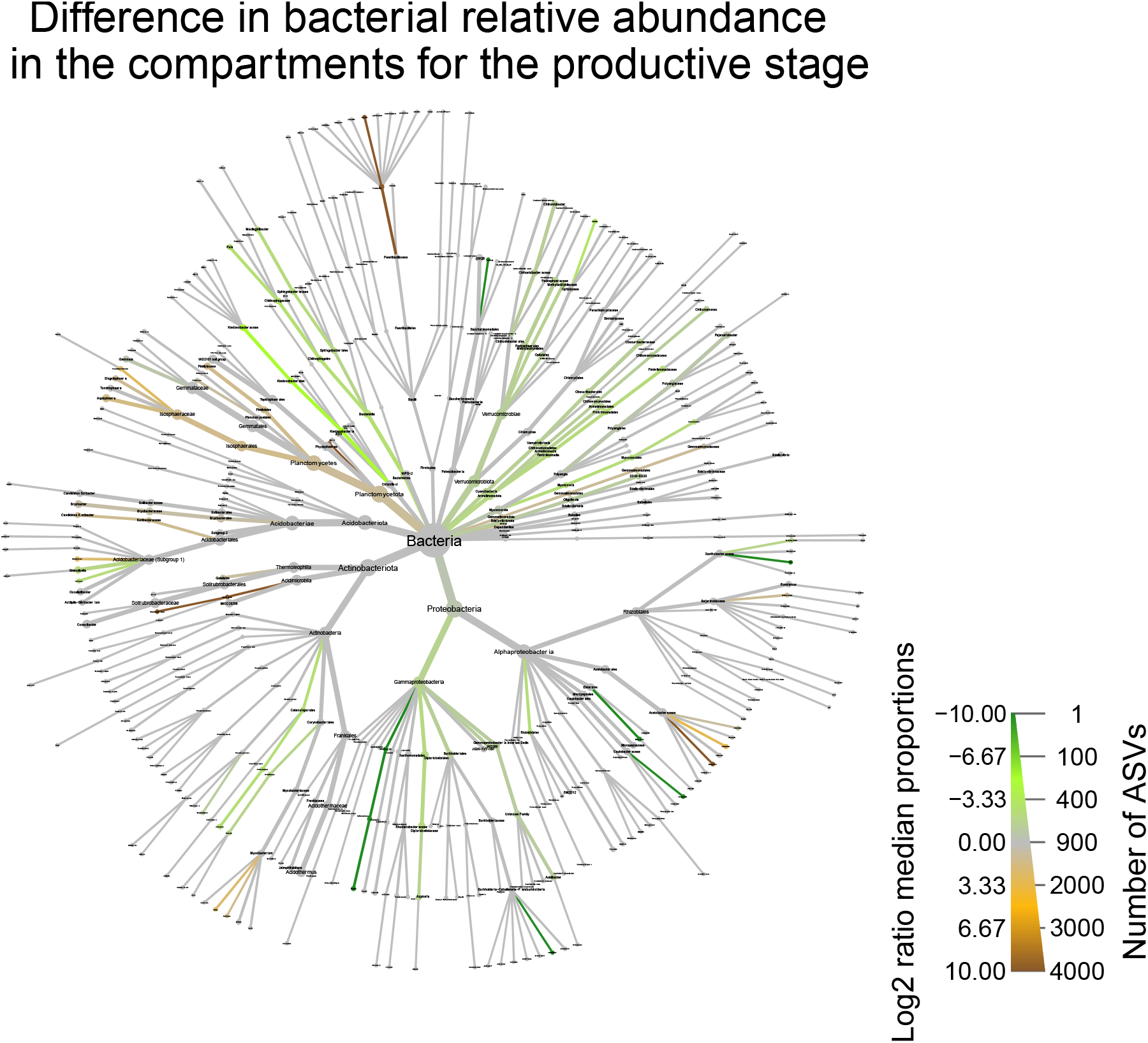
Differential abundance in the bacterial community for the harvesting year comparing the root and rhizosphere. The coloured ASVs have a significant difference in relative abundance in either plant compartment (green for roots, brown for rhizosphere). A log2ratio of 10 (dark brown) or −10 (dark green) indicates that the taxon was exclusively present in the given plant compartment. The size of the edges and nodes indicate the number of ASVs assigned to a particular taxonomic level.

**Supplementary Figure 12.**
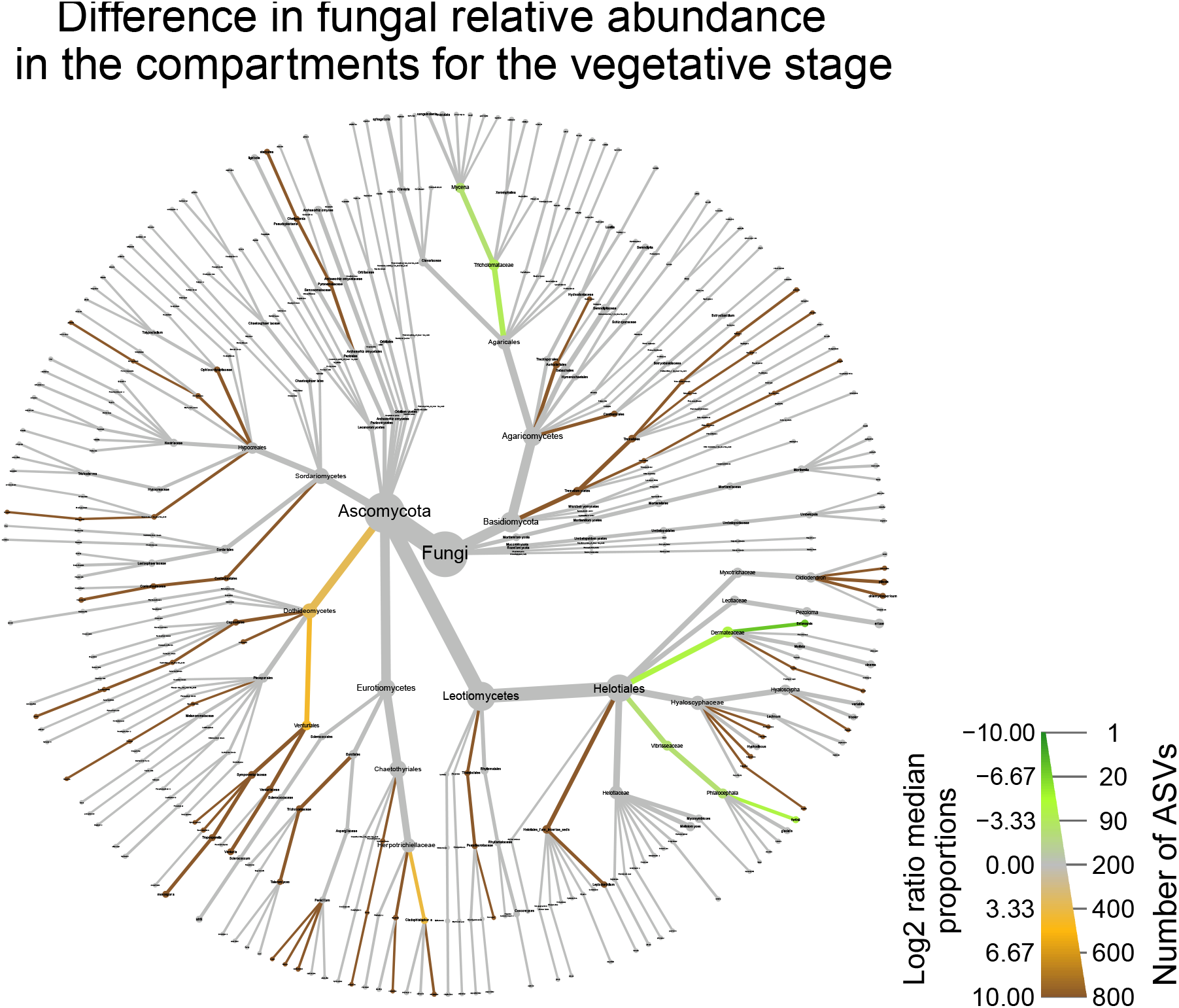
Differential abundance in the fungal community for the pruning year comparing the root and rhizosphere. The coloured ASVs have a significant difference in relative abundance in either plant compartment (green for roots, brown for rhizosphere). A log2ratio of 10 (dark brown) or −10 (dark green) indicates that the taxon was exclusively present in the given plant compartment. The size of the edges and nodes indicate the number of ASVs assigned to a particular taxonomic level.

**Supplementary Figure 13.**
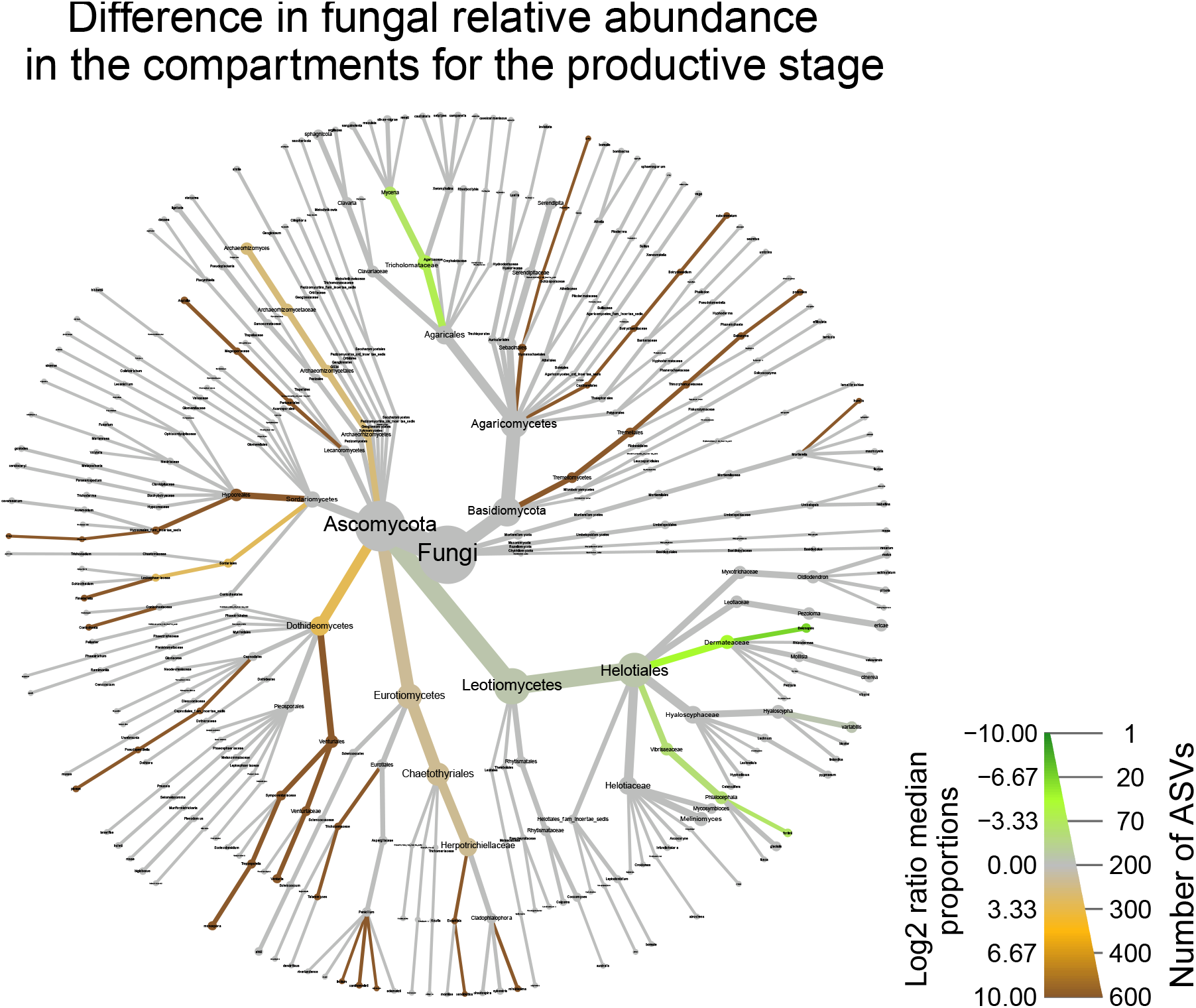
Differential abundance in the fungal community for the harvesting year comparing the root and rhizosphere. The coloured ASVs have a significant difference in relative abundance in either plant compartment (green for roots, brown for rhizosphere). A log2ratio of 10 (dark brown) or −10 (dark green) indicates that the taxon was exclusively present in the given plant compartment. The size of the edges and nodes indicate the number of ASVs assigned to a particular taxonomic level.

**Supplementary Table 1:** Bacterial mock community.

| Phylum | Order | Genus and species |
| --- | --- | --- |
| Proteobacteria | Pseudomonadales | *Acinetobacter baumannii* |
| Actinobacteriota | Actinomycetales | *Actinomyces odontolyticus* |
| Firmicutes | Bacillales | *Bacillus cereus* |
| Bacteroidota | Bacteroidales | *Bacteroides vulgatus* |
| Actinobacteriota | Clostridiales | *Clostridium beijerinckii* |
| Actinobacteriota | Propionibacteriales | *Propionibacterium acnes* |
| Deinococcota | Deinococcales | *Deinococcus radiodurans* |
| Firmicutes | Lactobacillales | *Enterococcus faecalis* |
| Proteobacteria | Enterobacterales | *Escherichia coli* K12 |
| Campylobacterota | Campylobacterales | *Helicobacter pylori* |
| Firmicutes | Lactobacillales | *Lactobacillus gasseri* |
| Firmicutes | Lactobacillales | *Listeria monocytogenes* |
| Proteobacteria | Burkholderiales | *Neisseria meningitidis* |
| Proteobacteria | Pseudomonadales | *Pseudomonas aeruginosa* |
| Proteobacteria | Rhodobacterales | *Rhodobacter sphaeroides* |
| Firmicutes | Staphylococcales | *Staphylococcus aureus* |
| Firmicutes | Staphylococcales | *Staphylococcus epidermidis* |
| Firmicutes | Lactobacillales | *Streptococcus pneumoniae* |
| Firmicutes | Lactobacillales | *Streptococcus agalactiae* |
| Firmicutes | Lactobacillales | *Streptococcus mutans* |

**Supplementary Table 2:** Effect of plant compartment and fertilization on bacterial and fungal communities using PERMANOVA test on Aitchison distance matrices.

| Factor | df | Pruning |  | Harvesting |  |
| --- | --- | --- | --- | --- | --- |
|  |  | Bacteria | Fungi | Bacteria | Fungi |
| R<sup>2</sup> • F value (p value) |  |  |  |  |  |
| Plant compartment (Pc) | 1 | 0.10 • **2.58 (0.001)** | 0.11 • **2.72 (0.001)** | 0.14 • **3.65 (0.001)** | 0.12 • **3.03 (0.001)** |
| Fertilization (Fe) | 2 | 0.11 • **1.42 (0.001)** | 0.11 • **1.29 (0.012)** | 0.10 • **1.33 (0.029)** | 0.11 • **1.34 (0.007)** |
| Pc x Fe | 2 | 0.06 • 0.73 (0.926) | 0.05 • 0.63 (0.997) | 0.06 • 0.76 (0.988) | 0.05 • 0.65 (0.999) |

Note : df, degree of freedom; significant results are in bold

